# Early alterations of RNA metabolism and splicing from adult corticospinal neurons in an ALS mouse model

**DOI:** 10.1101/667733

**Authors:** Christine Marques, Mathieu Fischer, Céline Keime, Thibaut Burg, Aurore Brunet, Jelena Scekic-Zahirovic, Caroline Rouaux

## Abstract

Amyotrophic lateral sclerosis (ALS) is a devastating neurodegenerative disease clinically defined as the combined degeneration of corticospinal and corticobulbar neurons (CSN), and bulbar and spinal motor neurons (MN). A growing body of evidence points to the motor cortex, where CSN are located, as the potential initiation site of ALS. However, little is known about the spatiotemporal dynamics of CSN degeneration and the molecular pathways involved. Here, we show in the *Sod1^G86R^* mouse model of ALS that CSN loss precedes MN degeneration and that CSN and MN degenerations are somatotopically related, highlighting the relevance of CSN to ALS onset and progression. To gain insights into the molecular mechanisms that selectively trigger CSN degeneration, we purified CSN from the motor and somatosensory cortex of adult mice and analysed their transcriptome from presymptomatic ages to disease end-stage. Significant RNA metabolism and splicing alterations, novel in the context of *Sod1* mutation, were identified, including mis-splicing events that largely trigger genes involved in neuronal functions. Together, the data indicate that CSN dysfunction and degeneration upon mutant *Sod1* expression involve alterations of RNA metabolism and splicing, emphasizing shared mechanisms across various ALS-related genes.

---

ALS is an incurable and fatal neurodegenerative disease that mostly starts in adulthood with a rapidly progressing paralysis and death within only 2 to 5 years of diagnosis^1,2^. In clinics, ALS is defined as the combined degeneration of both corticospinal and corticobulbar neurons (CSN, or upper motor neurons) whose cell bodies are located in the cerebral cortex and that extend axons to the medulla and spinal cord, and of spinal and bulbar motor neurons (MN, or lower motor neurons) whose cell bodies are located in the medulla and spinal cord, and that connect to the skeletal muscles. If the cellular and molecular mechanisms behind MN degeneration are relatively well known, the numerous related clinical trials have unfortunately failed to translate into improved treatment of ALS^1-4^, a discrepancy that contributed to the emergence of the CSN and the overall cerebral cortex as alternative therapeutic targets.

A growing body of evidence points to the motor cortex as the likely initiation site of ALS^5,6^. A first series of arguments arose from the recent genetic, pathological and clinical evolution links established between ALS and FrontoTemporal Dementia (FTD), a neurodegenerative disease that affects solely the cerebral cortex and results in behavioural and cognitive deficits^7^. A second series of arguments emerged following extensive examination of the TDP-43 aggregates, a pathological hallmark of the disease in post-mortem brains from ALS patients, identifying the motor cortex as the most affected region of the brain, and leading to the corticofugal hypothesis^8-10^ which proposes a sequential pattern of pathology spreading from the motor cortex to its projection sites. Finally, transcranial magnetic stimulation studies of ALS patients revealed typical hyperexcitability of the motor cortex that characterizes both sporadic and familial ALS patients, negatively correlates with survival, and manifests prior to disease onset^11^, further suggesting a primary involvement of the motor cortex in disease initiation. In this context, the CSN, that connect the motor cortex to its downstream targets and degenerate in ALS, appear as potentially very relevant cellular players of disease onset and progression.

So far, only a limited number of studies reported on the degeneration of CSN in mouse models of the disease. Early and progressive loss of CSN was initially demonstrated in the *SOD1^G93A/G1H^* and then the *SOD1^G93A^* mouse lines^12-14^. More recently, loss of CSN has been reported at symptom onset in the *Thy1::TDP43^A315T^* mice^15^. Finally *hPFN1^G118V^* mice^16^ and *C9-BAC* mice^17^ present respectively a loss of CTIP2-positive neurons or of large pyramidal cells within the layer V of the motor cortex, at disease end-stage. Together, these studies highly suggest that the mouse strains that best mimic ALS symptoms recapitulate CSN degeneration.

Here, we sought to unravel the signalling pathways that accompany CSN initial dysfunction and ultimate loss during the course of the disease, using the *Sod1^G86R^* mice that typically recapitulate ALS onset and progression, with first motor symptoms detected at around 90 days of age and premature death at around 110 days^18,19^. In the absence of specific reporter to label CSN, *i.e.* to discriminate them from the neighbouring and highly similar subcerebral projection neurons that populate the layer V of the motor and non-motor cerebral cortex, we performed series of retrograde labelling of this disease-relevant neuronal population. Rigorous sampling and counting method of CSN highlights their pre-symptomatic and progressive loss, and a somatotopic relationship between CSN and MN degenerations, which had not been previously reported in animal models. After purification of labelled CSN from the motor and somatosensory cortex of adult wild-type or *Sod1^G86R^* at two presymptomatic and two symptomatic ages, we conducted a longitudinal transcriptome analysis of this population under neurodegenerative conditions. We unravelled alterations of expression of genes involved in RNA metabolism, further confirmed by occurrences of RNA mis-splicing. These uncovered signalling pathways are highly reminiscent of those described for other, recently identified ALS-related genes known to regulate RNA metabolism, such as *FUS* and *TARDBP*^7^, but are novel for *SOD1*, highlighting common, disease-relevant molecular pathways.

## Results

### CSN progressively degenerate in *Sod1^G86R^* mice

To test whether *Sod1^G86R^* mice recapitulate a progressive loss of CSN, we performed a series of retrograde labelling of the whole CSN population from the cervical portion of the spinal cord of wild-type and *Sod1^G86R^* animals, by injecting Fluorogold at two presymptomatic ages, 60 days (60d: n=5 WT and 6 *Sod1^G86R^*) and 75 days (75d: n=6 WT and 3 *Sod1^G86R^*), and two symptomatic ages, 90 days (90d: n=5 WT and 6 *Sod1^G86R^*) and 105 days (105d: n=5 WT) or end-stage (ES: n=5 *Sod1^G86R^*) (Fig. 1a). Microscopy analysis of coronal sections revealed labelled pyramidal cells in the cerebral cortex, spanning from Bregma 2.10 mm to Bregma −2.30 mm (Supplementary Fig.1). As exemplified at the rostro-caudal level Bregma 0.74 mm, labelled cells located in cortical layer V span from the secondary motor cortex medially to the primary somatosensory cortex laterally, in both wild-type and *Sod1^G86R^* animals (Fig. 1b). We observed a marked decrease of labelled CSN in the cerebral cortex of *Sod1^G86R^* mice over time (Fig. 1b,c).

**Figure 1:**
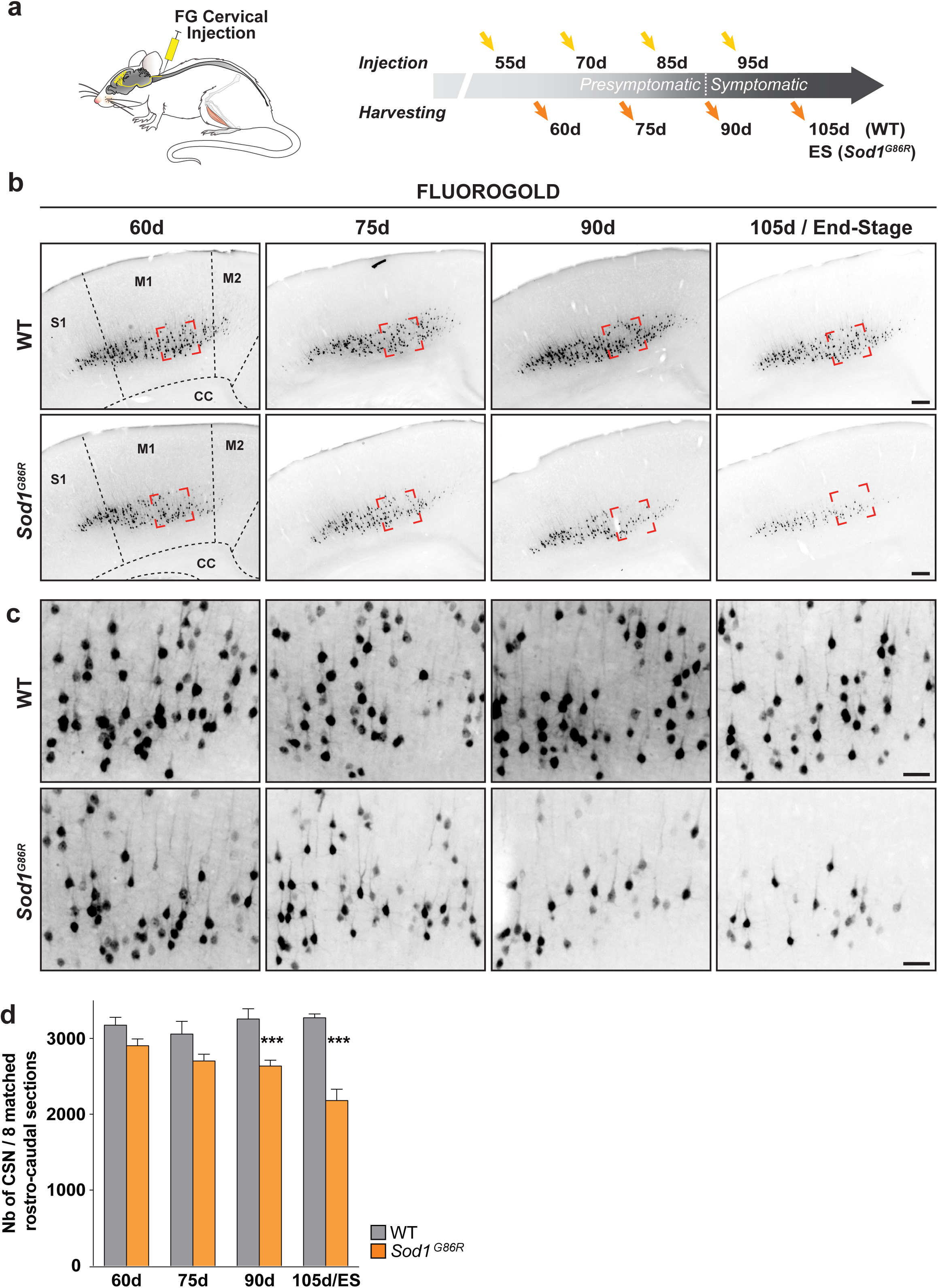
CSN progressively degenerate in *Sod1^G86R^* mice. **a.** Schematic of the experimental design: CSN were retrogradelly labelled upon Fluorogold (FG) injection into the cervical portion of the spinal cord. Ages of injection and harvesting are indicated. **b.** Representative negatives of fluorescence images of brain coronal sections (Bregma 0.74 mm) showing Fluorogold-labelled CSN in the cerebral cortex of wild-type and *Sod1^G86R^* mice of 60, 75, 90 and 105 days of age (d) or end stage (ES). Red shapes indicate the positions where close-ups were acquired. **c.** Close-ups showing the progressive loss of labelled CSN over time in the cerebral cortex of *Sod1^G86R^* mice (lower panels) compared to their wild-type littermates (upper panels). **d.** Bar graph representing the average number of CSN present on 8 equally spaced coronal sections along the rostro-caudal axis, matched between *Sod1^G86R^*(orange) and wild-type mice (grey). Note the progressive loss of labelled CSN in the brain of *Sod1^G86R^* animals. *** p<0.001 in multiple t tests. Scale bars: 200 µm in b. and 50 µm in c.

To evaluate a potential gradual loss of the whole population of CSN in *Sod1^G86R^* animals, we quantified the number of labelled CSN present in both hemispheres of 8 coronal sections equally spaced along the rostro-caudal axis and matched between the two genotypes (Supplementary Fig. 1). The results show that the population of retrogradelly-labelled CSN progressively decreased over time in *Sod1^G86R^* animals compared to their wild-type littermates (Fig. 1d). This decrease became significant at the symptomatic age of 90 days, and reached a loss of 33.3% by the end of the disease (Fig. 1d).

To confirm these results and exclude the possibility that the loss of CSN could be due to a defective retrograde transport of Fluorogold by the axons of the *Sod1^G86R^* CSN, we took advantage of the expression of the *mu-Crystallin* gene (*Crym*) by the CSN, along with other subcerebral projection neurons of the cortical layer V^20^. *In situ* hybridization for *Crym* (Supplementary Fig. 2), was performed at one single rostro-caudal level (Bregma 0,14 mm) and within the layer V of the secondary and primary motor area and the most medial part of the primary sensorimotor areas only, at 60d (n=3 WT and 3 *Sod1^G86R^*), 90d (n=4 WT and 4 *Sod1^G86R^*), 105d (n=4 WT and 4 *Sod1^G86R^*) and 115d/ES (n=3 WT and 3 *Sod1^G86R^*). Quantification revealed a progressive reduction of *Crym*-positive neurons in *Sod1^G86R^* mice compared to their wild-type littermates (Supplementary Fig. 2). While the data do not rule out the possibility that retrograde transport may be altered in CSN over time, they nevertheless confirm a progressive loss of the CSN population in *Sod1^G86R^*.

### CSN degeneration propagates in a caudo-rostral manner in *Sod1^G86R^* mice

In mice, CSN are topographically organized along the rostro-caudal and the medio-lateral axes, and project accordingly to different portions of the spinal cord^21,22^. Sampling the whole population of CSN along the rostro-caudal axis (Supplementary Fig. 1) enabled us to test whether their degeneration was homogeneous, or rather affected more selectively sub-populations. We thus quantified the number of CSN present on each of the 8 selected coronal sections (Fig. 2) of wild-type and *Sod1^G86R^* brains over time. Using this approach, we observed that the number of CSN was uneven along the rostro-caudal axis, and that CSN were overall more numerous in the most caudal sections (Fig. 2 and Supplementary Fig.1). At the presymptomatic ages 60d and 75d, *Sod1^G86R^* brains displayed less CSN than their wild-type counterparts, at every level analysed, but no significant difference could be detected between the two genotypes (Fig. 2). At 90d, we observed a significant reduction of CSN in the caudal sections only, and this reduction was uneven amongst these sections, with a greater decrease observed at the most caudal levels (section 5: 11,86% loss; 6: 19,23%; 7: 19,21%; 8: 29,59%; 9: 28,43%; Fig. 2). At 105d/ES, with the exception of section 3, the loss of CSN was significant at every level. Within the caudal part, section 9 remained the most severely affected, with a loss of 44,48%, but overall, the most impacted section was the rostral section 2, with a loss of 68,67% (Fig. 2). Together, the data show that CSN degeneration is not even along the rostro-caudal axis in *Sod1^G86R^* mice: it starts with the most caudally located ones, and progresses rostrally over time.

**Figure 2:**
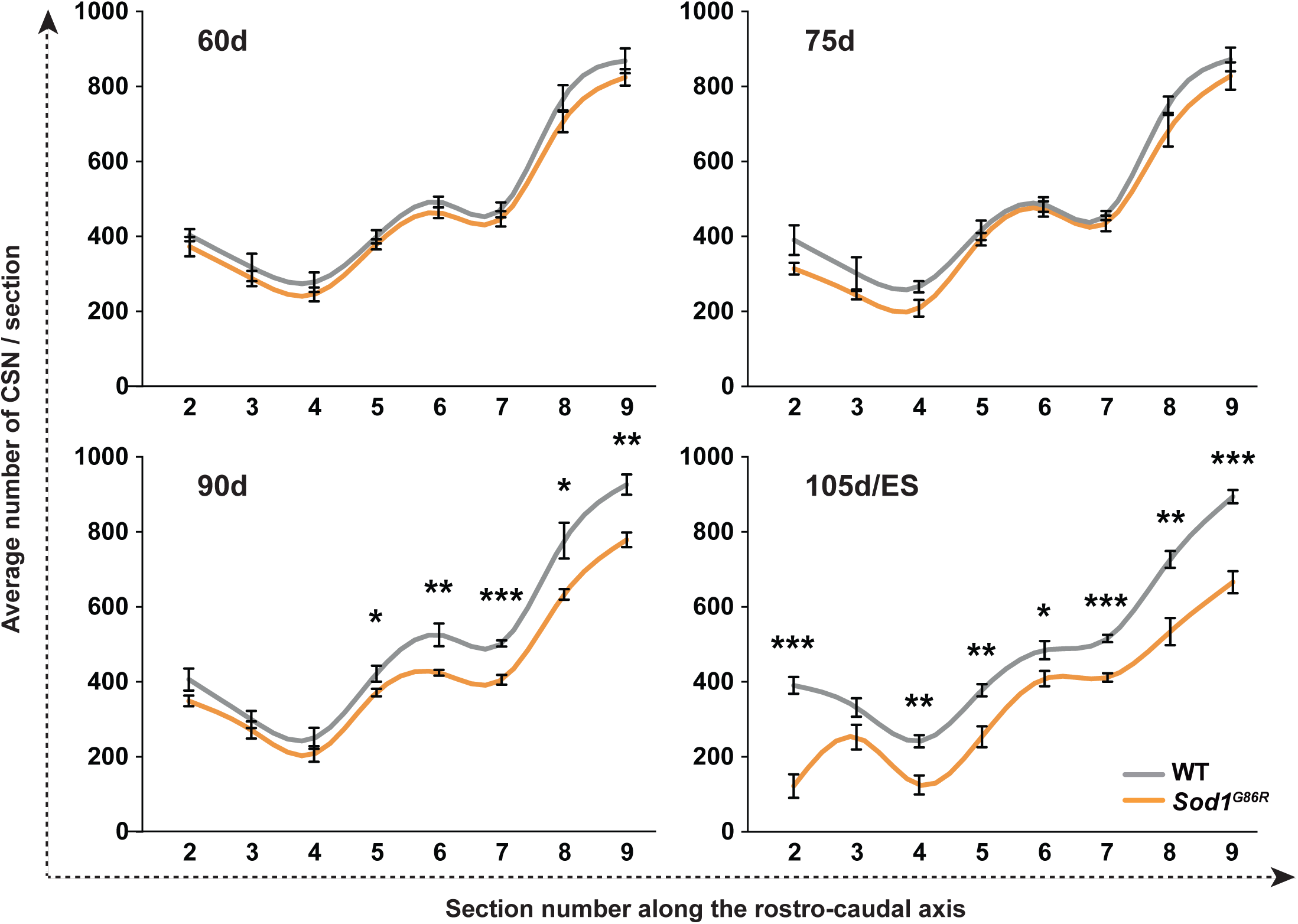
In *Sod1^G86R^* mice, CSN degeneration starts caudally and progresses rostrally. Graphs representing the average number of Fluorogold-labelled CSN counted on both hemispheres of equally spaced coronal sections 2 (Bregma 2.10 mm) to 9 (Bregma −1.70 mm) of *Sod1^G86R^* mice (orange) and their wild-type littermates (grey) at 60, 75, 90 and 105 days of age (d) or end stage (ES). * p<0.05; ** p<0.01; *** p<0.001 in multiple t-tests.

### CSN and MN degenerations are temporally and somatotopically related

Because CSN that innervate the lumbar spinal cord are mostly caudally located^21,22^, we reasoned that they might account for the majority of the neuronal loss observed in the most caudal sections we analysed. To test this hypothesis, we performed a second set of retrograde labelling and injected Fluorogold from the lumbar portion of the spinal cord of wild-type and *Sod1^G86R^* animals, at the same presymptomatic ages (60d: n=4 WT and 3 *Sod1^G86R^*; 75d: n=4 WT and 3 *Sod1^G86R^*) and symptomatic ages (90d: n=5 WT and 5 *Sod1^G86R^*; 105d: n=4 WT and ES: n=4 *Sod1^G86R^*) (Fig. 3a). Microscopy analysis of coronal sections revealed labelled cortical cells, spanning from Bregma −0,46 mm to Bregma −1,70 mm (Supplementary Fig. 3). Each of the three levels considered (section 8, Bregma −1,06 mm, Fig.3, and sections 7 and 9, Bregma −0,46 and −1,70 mm respectively, data not shown), showed a marked loss of CSN from 60 days on (Fig. 3c). Sampling and counting the lumbar-projecting CSN revealed a marked loss of this subpopulation in the *Sod1^G86R^* mice compared to their wild-type littermates, which was significant from 60 days on, and intensified over time (Fig. 3b). Taken together, the data show that, in the *Sod1^G86R^* animals, the loss of lumbar-projecting CSN starts earlier than that of the overall CSN population.

**Figure 3:**
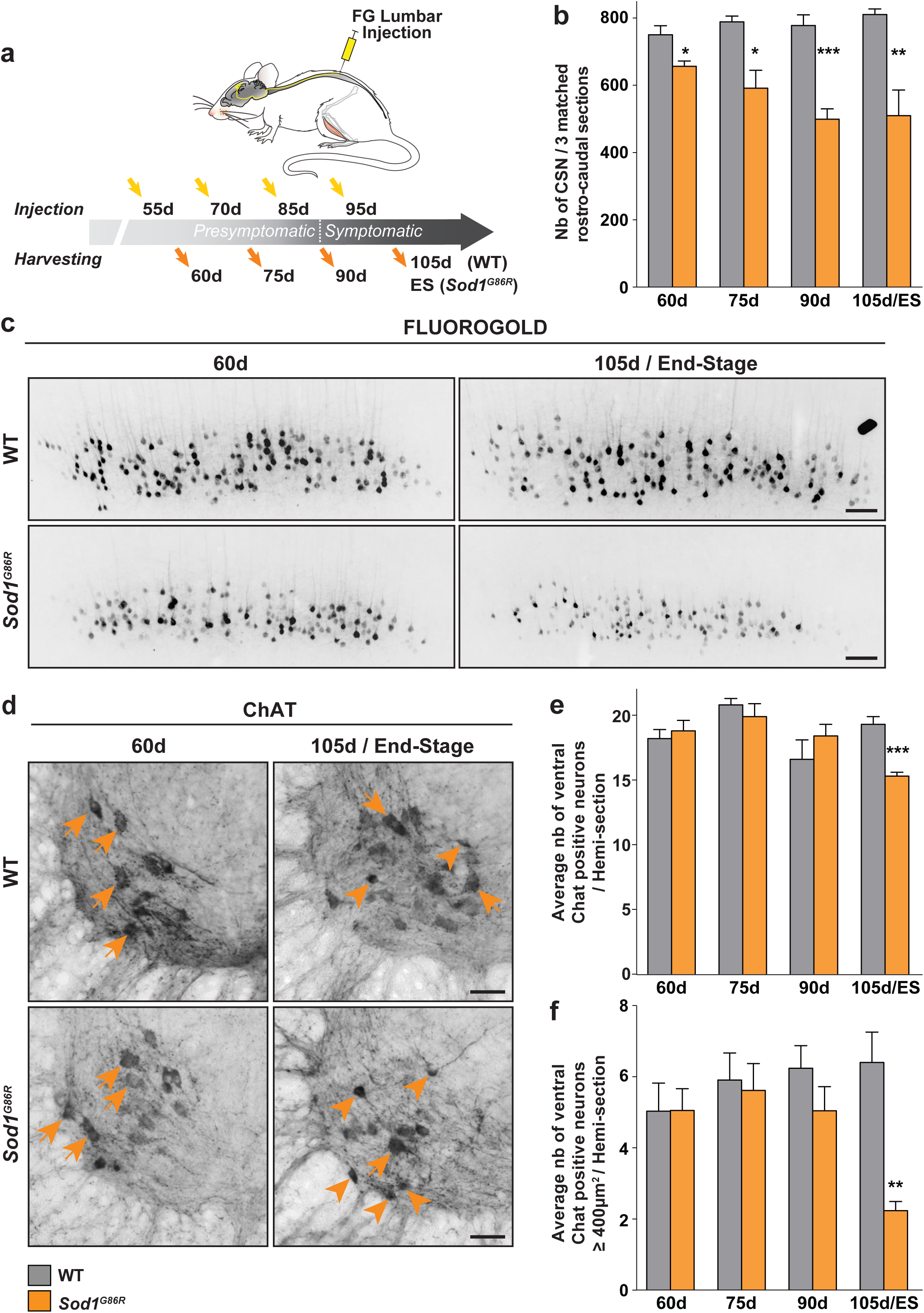
Lumbar-projecting CSN degenerate before lumbar MN. **a.** Schematic of the experimental design: CSN were retrogradelly labelled upon Fluorogold (FG) injection into the lumbar portion of the spinal cord. Ages of injection and harvesting are indicated. **b.** Bars graph representing the average number of CSN present on 3 equally spaced caudal coronal sections matched between *Sod1^G86R^* (orange) and wild-type mice (grey), at 60, 75, 90 and 105 days of age (d) or at end stage (ES). Note the early and progressive loss of labelled CSN in the brain of *Sod1^G86R^* animals. **c.** Representative negatives of fluorescence images of brain coronal sections (Bregma −1.06 mm) showing the decreased number of Fluorogold-labelled CSN in the cerebral cortex of *Sod1^G86R^* mice at 60 days of age and at end stage (ES) compared to their wild-type littermates. **d.** Representative images of coronal sections of the spinal cord of wild-type and *Sod1^G86R^* mice at 60 and 105 days (d) or end stage (ES), showing Choline AcetylTransferase-positive neurons (ChAT) present in the ventral horn of the spinal cord. Arrows indicate large ChAT-positive neurons, arrowheads indicate smaller or shrinked ChAT-positive neurons. **e.** Bar graph representing the average number of ChAT-positive neurons counted in the ventral horn of lumbar spinal cord sections of wild-type (grey) and *Sod1^G86R^* mice (orange) at 60, 75, 90 and 105 days of age end stage (ES). **f.** Bar graph representing the average number of large ChAT-positive neurons, with an area ≥ 400 µm^2^, counted in the ventral horn of lumbar spinal cord sections from wild-type (grey) and *Sod1^G86R^* mice (orange) at 60, 75, 90 and 105 days of age or end stage (ES). * p<0.05; ** p<0.01; *** p<0.001 in multiple t-tests. Scale bars: 100 µm in c. and 50 µm in d.

The paralysis that progressively affects the *Sod1^G86R^* mice typically starts in the hind limbs. In accordance with this, spinal motor neurons (MN) were shown to degenerate in the lumbar portion of the spinal cord^18,19^. In order to compare the time course of lumbar-projecting CSN degeneration to that of lumbar MN, we counted the number of ChAT-positive neurons present in the ventral horn of the lumbar spinal cord of wild-type and *Sod1^G86R^* animals over time (60d: n=5 WT and 6 *Sod1^G86R^*; 75d: n=6 WT 3 *Sod1^G86R^*; 90d: n=5 WT and 6 *Sod1^G86R^*; 105d/ES: n=5 WT 5 *Sod1^G86R^*; Fig. 3d,e,f). We failed to detect any significant difference between *Sod1^G86R^* and wild-type mice before end-stage of the disease (15,3 ± 0,3 vs 19,3 ± 0,6 respectively, corresponding to a loss of 20,7%; Fig. 3e). To exclude the contribution of small, disease-resistant gamma motor neurons and restrict the counting to large, disease-vulnerable alpha motor neurons, we next counted the number of large (400µm^2^ or above) ventral ChAT-positive neurons (Fig. 3f). We observed an important and significant loss of large ChAT-positive neurons at end-stage of the disease (65,27%, p<0,001), and a milder and non-significant loss at 90 days of age (19,19%, p=0,237). Together, the data suggest that, in *Sod1^G86R^* mice, actual loss of lumbar MN following the shrinkage of their somas is a rather late event in comparison with the overall disease progression.

In mutant *Sod1* mouse models of ALS, MN undergo Wallerian degeneration, characterized by initial alteration of the neuromuscular junction (NMJ) and retrograde progression to the soma^23^. To determine when lumbar MN degeneration starts, we performed electromyography recordings in the tibialis anterior (Supplementary Fig. 4a,b) muscles of both hind limbs of the *Sod1^G86R^* mice and their control littermates prior to their harvesting. Using this technique, NMJ denervation was evident on 100% of *Sod1^G86R^* end-stage animals, and 66,66% of 90 day-old animals, but not on younger animals (Supplementary Fig. 4a,b). To confirm these results, we investigated the status of the NMJ of the tibialis anterior muscle by immunofluorescence (Supplementary Fig. 4c,d). In both wild-type and *Sod1^G86R^* animals we observed healthy, innervated neuromuscular junctions, along with a small proportion of partly denervated NMJ that displayed discrete disconnections of the axonal terminal (Supplementary Fig.4d). The proportion of partly denervated NMJ was stable across ages and genotypes (Supplementary Fig.4c), indicating that their occurrence likely reflects a mild alteration of the tissue during the process of bundles preparation. Importantly, in *Sod1^G86R^* animals only, we detected fully denervated NMJ that had lost their axonal terminal (Supplementary Fig.4d). This phenotype was present only from 90 days on, and progressed over time (17,7% of *Sod1^G86R^* mice at 90 days, 76,9% at end-stage, Supplementary Fig. 4c). Thus, the data indicate that lumbar MN degeneration starts after 75 days of age in *Sod1^G86R^* mice.

Taken together, the data show that in a mouse model of ALS where paralysis starts in the hind limbs, CSN degeneration occurs in a somatotopic manner, with lumbar-projecting CSN being affected earlier than the rest of the CSN population. In addition, the data indicate that CSN loss occurs before the appearance of the first signs of MN degeneration. The spatiotemporal dynamics of CSN degeneration during the course of the disease in *Sod1^G86R^* mice suggests that CSN are likely affected before the MN, further highlighting their clinical relevance.

### Purification of adult CSN under healthy and neurodegenerative conditions

To investigate the molecular mechanisms behind CSN degeneration, we first developed an approach to purify this neuronal population from the cerebral cortex of adult wild-type and *Sod1^G86R^* mice. CSN were retrogradelly labelled by injecting green fluorescent microspheres into the cervical portion of the dorsal funiculus at 25 days of age (Fig. 4a,b,c). Animals were harvested at two presymptomatic ages, 30 and 60 days, and two symptomatic ages, 90 and 105 days (Fig. 4b). CSN-containing cortical layers V were then microdissected and labelled CSN were dissociated and purified by FACS (Fig. 4a,c). In parallel, a second population of cortical neurons, the callosal projection neurons (CPN) located in the layers II/III of the cerebral cortex, above the labelled CSN, were also purified (Fig. 4a,c). CPN and CSN are close neuronal populations: they arise from a same pool of cortical progenitors, are excitatory projection neurons that use the glutamate as a neurotransmitter, and express some common molecular markers. Yet, as opposed to CSN, CPN have not been reported to degenerate in ALS, and may thus represent a valuable control population of cortical excitatory neurons that overexpress the *Sod1^G86R^* transgene without degenerating. To confirm this, we took advantage of *Cux2* selective expression in upper layer CPN^24^, and measured the density of CUX2-positive nuclei in the cortical layers II/III of wild-type and *Sod1^G86R^* at 60, 75, 90 and 105 days of age or at disease end-stage. No decrease of CUX2-positive nuclei could be detected (Supplementary Fig. 5). Instead, we observed a significant increase of CUX2 density at disease end stage, which could potentially arise from an overall decrease of the cortical thickness. We reasoned that the comparison between the two neuronal populations, vulnerable CSN versus resistant CPN, would enable to distinguish genes dysregulated in CSN only, and likely to be involved in their degeneration, from genes dysregulated in both populations, and unlikely to have any detrimental effect on neuronal survival. Importantly, in most of the cases, CSN and CPN were purified from same animals (Fig. 4a,c). 64 samples were initially sequenced (n=4 biological replicated × 2 genotypes × 2 cell types × 4 ages), 7 were excluded because they did not expressed the expected markers or were altered, and a total of 57 samples were further processed (CSN: 30d: n=4 WT and 3 *Sod1^G86R^*; 60d: n=4 WT and 4 *Sod1^G86R^*; 90d: n=4 WT and 4 *Sod1^G86R^*; 105d: n=4 WT and 2 *Sod1^G86R^*; CPN: 30d: n=4 WT and 3 *Sod1^G86R^*; 60d: n=2 WT and 3 *Sod1^G86R^*; 90d: n=4 WT and 4 *Sod1^G86R^*; 105d: n=4 WT and 4 *Sod1^G86R^*). Unsupervised hierarchical clustering of 45,337 mRNA transcripts in all samples clearly separated the two neuronal populations in distinct clusters (Fig. 4d). qPCR (data not shown) and RNAseq data (Fig. 4e) confirmed that purified CSN expressed markers typical of the cortical layer V subcerebral projection neurons, and did not express any marker typical of upper layer projection neurons^20,24,25^, and vice-versa (Fig. 4e). Together, the data indicate that subtypes of excitatory neuronal populations can be successfully co-purified from individual adult mouse cortices, under healthy or neurodegenerative conditions.

**Figure 4:**
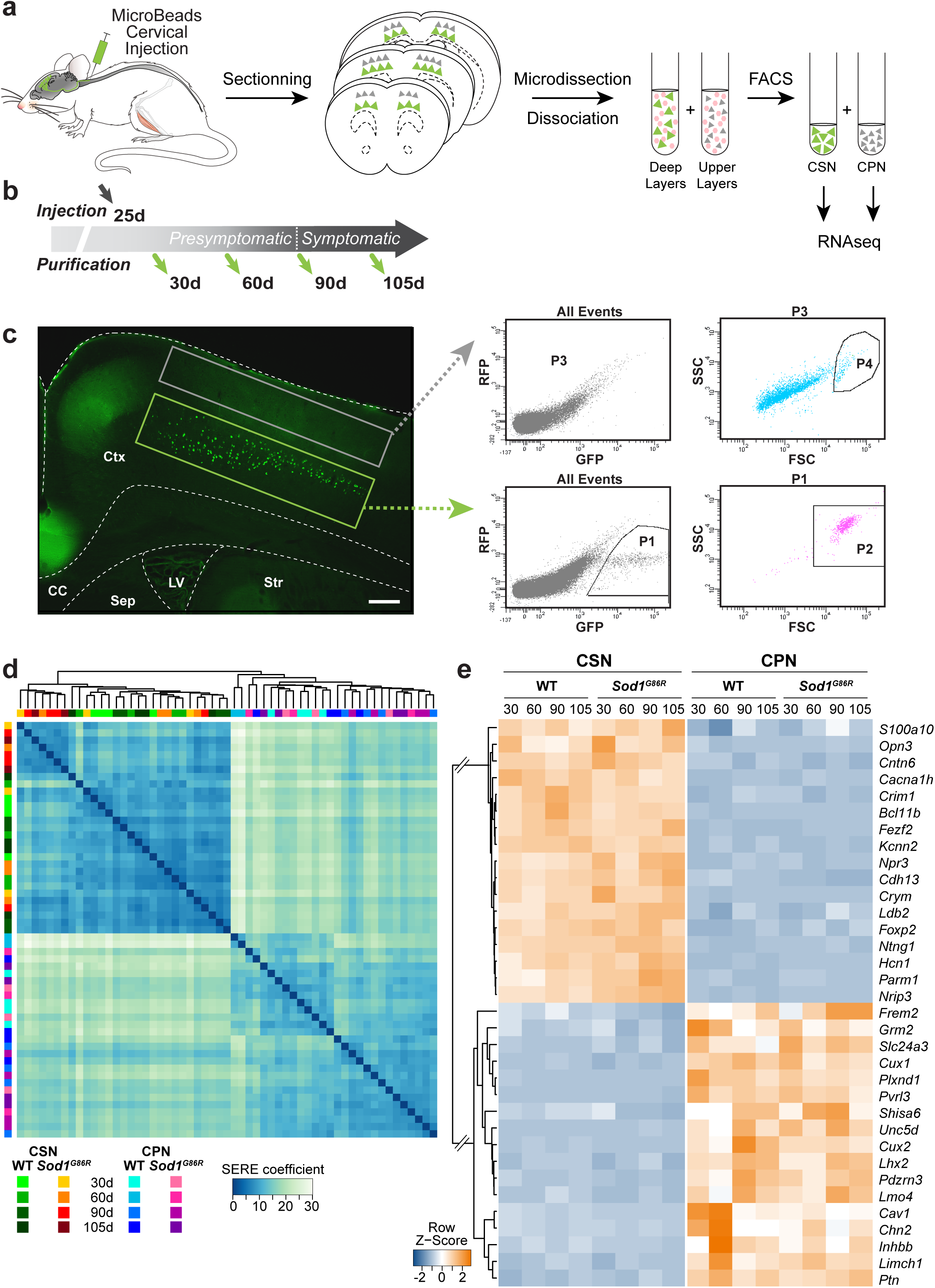
Isolation of CSN and CPN from individual adult wild-type and *Sod1^G86R^* mouse brains for RNAseq time course analysis. **a.** Schematic representation of the experimental design: CSN were retrogradelly labelled upon green fluorescent microbeads injection into the cervical portion of the spinal cord. The brains were then sectioned coronally to allow the microdissection of the layers II/III and V separately, followed by dissociation of the tissues and FACS purification of the CSN and CPN populations. **b.** Schematic representation of the ages of CSN labellings, and CSN and CPN harvestings. **c.** Representative fluorescence image of a brain coronal section upon CSN retrograde labelling, showing green fluorescent microbeads-labelled CSN within the layer V of the motor and somatosensory areas, and indicating the two microdissected areas (left, green and grey boxes; scale bar: 200 µm), along with representative FACS plots (right) of CPN (top) and CSN (bottom) purifications. RFP: Red Fluorescent protein; GFP: Green Fluorescent Portein; SSC: Side scatter; FSC: Forward scatter. **d.** Heatmap with unsupervised clustering of 45,337 mRNA transcripts in 57 samples calculated on the SERE coefficient that quantifies global RNA-seq sample differences. Shades of green or red identify wild-type versus *Sod1^G86R^* CSN samples respectively across ages. Shades of blue or pink to identify wild-type versus *Sod1^G86R^* CPN samples respectively across ages. **e.** Heatmap of gene expression patterns for known subtype-specific genes confirms the identities of the two isolated neuronal populations, across genotypes and ages.

### RNAseq analysis reveals early, CSN-specific transcriptional alterations in *Sod1^G86R^* mice

When performed separately on CSN or CPN samples, PCA globally distinguished the wild-type from the *Sod1^G86R^* CSN (Fig. 5a, left panel), while no sub-groups were visible amongst the CPN samples (Fig. 5a, right panel). Taking all ages into account, analysis of differential gene expression between wild-type and *Sod1^G86R^* samples indicated that numerous genes present a statistically significant differential expression within the CSN samples (Fig. 5b, left panel), as opposed to the CPN samples (Fig. 5a, right panel), in spite of similar strong up-regulation of the *Sod1* mRNA levels in both populations (Fig. 5a,b, inserts). Next, we evaluated age by age, the proportion and significance of differentially expressed genes between wild-type and *Sod1^G86R^* CSN samples. As illustrated by the volcano plots (Fig. 5c), the number of genes significantly dysregulated in *Sod1^G86R^* CSN are more important in symptomatic than in pre-symptomatic ages. Finally, we compared the pools of genes significantly dysregulated between wild-type and *Sod1^G86R^* CSN samples, and between wild-type and *Sod1^G86R^* CPN samples (Fig. 5d). Venn diagrams at four ages of interest (Fig. 5d) illustrate that important transcriptional dysregulations selectively occurred in CSN. These selective dysregulations started as early as 30 days of age and became more numerous as neurodegeneration progressed. In comparison, CPN transcriptional dysregulations are very modest. Importantly, very few common genes were found between the two neuronal populations. Thus, the data suggest that the two neuronal populations cope differently with the overexpression of the mutant *Sod1^G86R^* transgene, in accordance with the selective and progressive degeneration of the CSN and the maintenance of the CPN in the *Sod1^G86R^* animals over time.

**Figure 5:**
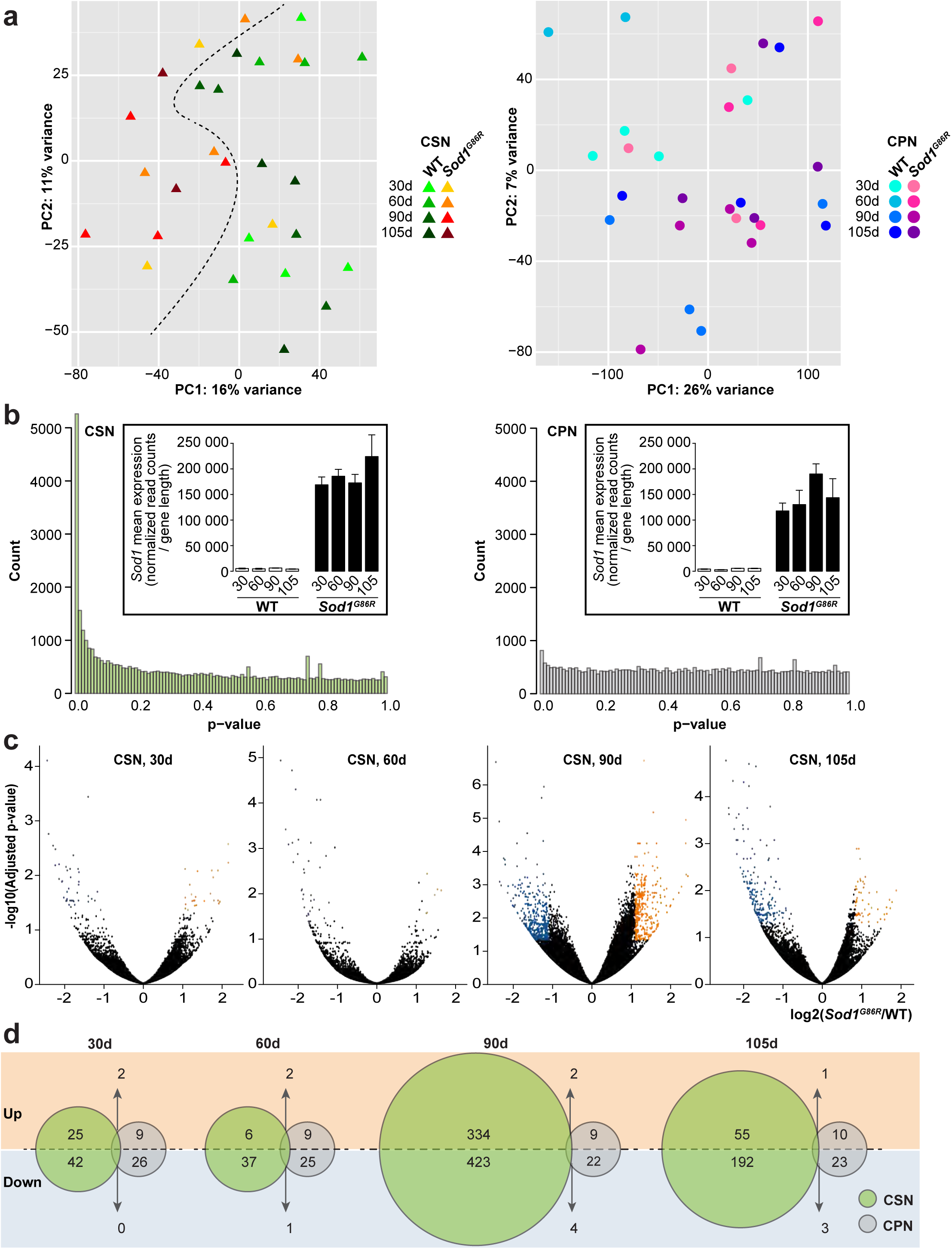
Disease progression differentially affects the transcriptomes of CSN and CPN in *Sod1^G86R^* mice. **a.** Principal component analyses ran on 45,337 mRNA transcripts in *n* = 29 CSN (left) and 28 CPN samples (right). Triangles represent CSN, with shades of green or red to identify wild-type versus *Sod1^G86R^* samples respectively. Circles represent CPN with shades of blue or pink to identify wild-type versus *Sod1^G86R^* samples respectively across ages. The dotted line highlights the segregation of *Sod1^G86R^* versus wild-type CSN. **b.** p-values histograms of CSN (green, left) versus CPN samples (grey, right), showing the large proportion of significantly dysregulated genes amongst CSN samples as opposed to CPN samples. Inserts represent the mean expression over replicate samples of *Sod1* mRNA across ages and genotypes for CSN (left) and CPN (right). **c.** Volcano plots of t-statistics, obtained upon exclusion of the overexpressed *Sod1* gene, showing the differential expression of genes between *Sod1^G86R^* and wild-type CSN at 30, 60, 90 and 105 days. For a better visualisation of these plots, *Sod1* and *Gm8566* genes have not been represented. Orange and blue dots respectively correspond to significantly up- and down-regulated genes. **d.** Venn diagrams showing the relative numbers of significantly dysregulated genes in the two neuronal populations, CSN in green and CPN in grey, at each age analysed. Upper halves of the circles over the orange background correspond to up-regulated genes, lower halves over the blue background to down-regulated ones.

### Gene set enrichment and network analyses resolve RNA metabolism dysregulation in *Sod1^G86R^* CSN

To identify the molecular pathways that accompany CSN degeneration, we ran a Gene Set Enrichment Analysis^26^ (GSEA) on a ranked list of 29, 017 genes significantly and selectively dysregulated in CSN and not in CPN across ages (Fig. 6a, and Supplementary Table 1). As expected in a context of ALS, the first gene ontology (GO) terms for biological processes (BP), cellular components (CC) and molecular functions (MF) were linked to the mitochondria (Fig. 6a, and Supplementary Table 1), indicating that CSN degeneration, like MN degeneration, is accompanied with alteration of the mitochondrial functions. Interestingly, amongst the 24 most up-regulated GO terms for BP, 7 were associated with RNA metabolism and transcriptional initiation: Ribonucleoprotein complex biogenesis and assembly (Normalized Enrichment Score, NES=2.94; Size=66), Protein RNA complex assembly (NES=2.86; Size=51), Translational initiation (NES=2.85; Size=34), RNA processing (NES=2.77; Size=136), tRNA metabolic process (NES=2.75; Size=16), Regulation of Translation Initiation (NES=2.63; Size=27) and RNA splicing (NES=2,62; Size=66) (Fig. 6a, and Supplementary Table 1). This relatively important representation of the GO terms in relation with RNA metabolism was further emphasized on an Enrichment Map for the BP, where the terms in relation with RNA metabolism appeared as part of the first sub-network (Fig. 6b, and Supplementary Fig. 6). In accordance with this, GO for CC indicated up-regulations of the Ribonucleoprotein complex (NES=3.59; Size=117 genes), the Endoplasmic reticulum (NES=3.20; Size=224 genes) and the Spliceosome (NES=2.65; Size=40) (Supplementary Table 1). Similarly, GO for MF indicated up-regulations of the Translation Initiation Factor Activity (NES=2.63; Size=20 genes), the Translation Regulator Activity (NES=2.38; Size=33 genes) and the Translation Factor Activity / Nucleic Acid Binding (NES=2.35; Size=31) (Supplementary Table 1). Expression profiles of some of these genes involved in RNA metabolism are shown for the pre-symptomatic and symptomatic ages of 30 and 90 days respectively (Fig. 6c). Remarkably, many of these genes are related either to motoneuron diseases such as Hereditary Spastic Paraplegia, Primary Lateral Sclerosis, Charcot-Marie-Tooth Disease, Spinal Muscular Atrophy, or to mental retardation and other related neurodevelopmental diseases, or both (Fig. 6c, asterisks, and Supplementary Table 2). We thus reasoned that some of these transcriptional alterations might occur very early on during the development and specification of the CSN. To test this possibility, we used microfluidics to run high-throughput qPCR validations on two groups of samples: cortical layers V microdissected from the motor cortex of 60 day-old wild-type and *Sod1^G86R^* animals, where the contribution of CSN to the overall variety of cell types is modest (Fig. 6d), and cortical plates from embryonic day (E) 14.5 wild-type and *Sod1^G86R^* embryos, that are enriched in developing layers V and VI corticofugal neurons and include specifying CSN, and are mostly devoid of other cortical excitatory projection neurons, cortical inhibitory neurons, and glial cell types (Fig. 6e). High-throughput qPCR results indicated that several of the gene expression dysregulations identified in pure adult CSN populations could also be observed in whole adult cortical layer V (Fig. 6d), in spite of the large dilution of the CSN contribution to the overall samples, and to a larger extent in E14.5 cortical plates of *Sod1^G86R^* embryos in comparison with their wild-type littermates (Fig. 6e). Together, the data indicate that CSN degeneration in *Sod1^G86R^* mice correlates with transcriptional alterations of genes involved in RNA metabolism. Interestingly, part of these transcriptional dysregulations are already present during corticogenesis, suggesting that CSN ultimate degeneration in the *Sod1^G86R^* mouse model of ALS could arise, at least partly, from neurodevelopmental alterations that may affects their proper specification, integration into neuronal circuits and functioning.

**Figure 6:**
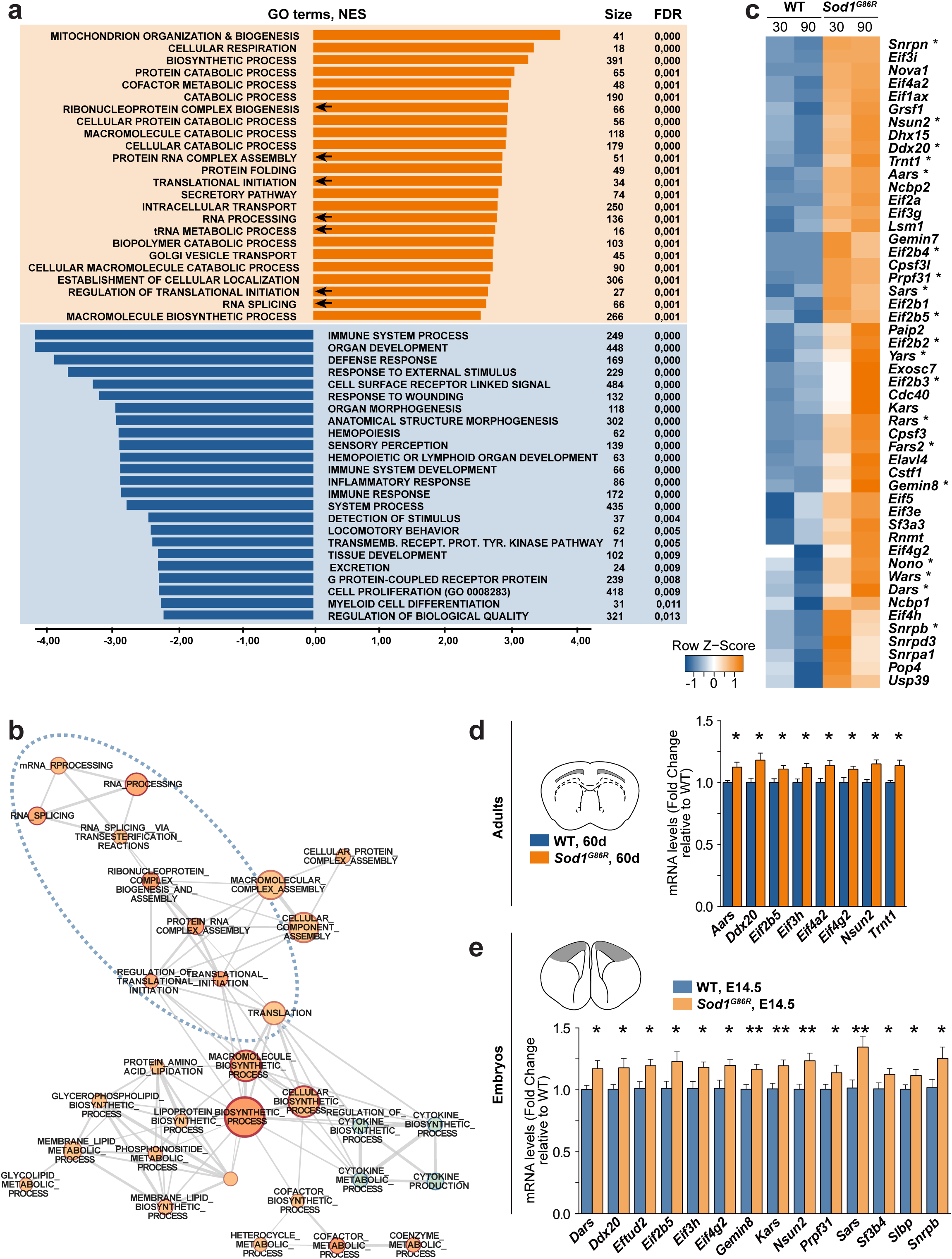
Gene Set Enrichment Analysis of diseased CSN reveals altered RNA metabolism. **a.** Gene set enrichment analysis (GSEA) of 29,017 ranked genes for GO terms of Biological Processes. The 15 most enriched positive (orange) and negative (blue) categories are listed according to their normalized enrichment score (NES) and False Discovery Rate (FDR), and the numbers of corresponding genes (Size) are indicated. Arrows indicate the redundancy and overrepresentation of GO terms related to RNA metabolism. **b.** Enrichment map representing the first sub-network identified with cytoskape. Nodes represent gene sets, and their size is proportional to the number of corresponding genes. Edges represent overlaps between gene sets, and their width is proportional to the number of overlapping genes. Node colour and intensity vary with the corresponding NES: orange for positive and blue for negative NES respectively, the darker the higher. Dotted blue line circles the GO terms linked to RNA metabolism. **c.** Heatmap of expression patterns of genes involved in RNA metabolism and identified by gene set enrichment analysis. Asterisks indicate genes related to motoneuron and/or neurodevelopmental diseases. **d.** Schematic representation of the microdissected layers V from 60 day-old wild-type and *Sod1^G86R^* mouse brains (left) and validation of candidate genes by high throughput qPCR (graph bar, right). **d.** Schematic representation of the microdissected cortical plate from E14.5 wild-type and *Sod1^G86R^* embryos (left) and validation of candidate genes by high throughput qPCR (graph bar, right). * p<0.05, ** p<0.01 between wild-type and *Sod1^G86R^* samples.

### RNA mis-splicing accompanies CSN degeneration

The identification of broad transcriptional alterations of genes involved in RNA metabolism, and more particularly in RNA splicing, may translate into altered mRNA splicing profile of the *Sod1^G86R^* CSN compared to the wild-type CSN population. To test this possibility, we employed rMATS^27^ to detect alternative splicing events from the 90 day-old wild-type and *Sod1^G86R^* CSN (n=4 per group) and CPN (n=2 WT and 3 *Sod1^G86R^*) replicates (Fig. 7). A total of 1,707 CSN-selective alternative splicing events corresponding to 1,163 affected genes were identified as skipped exons (SE, 722 events, 42.3%), alternative 5’ splice sites (A5SS, 241 events, 14.1%), alternative 3’ splice sites (A3SS, 334 events, 19.6%), mutually exclusive exons (MXE, 224 events, 13.1%) and retained introns (RI, 186 events, 10.9%) (Fig. 7a). GSEA analysis ran on the affected genes revealed the overrepresentation of GO terms for biological processes related to Neuronal Differentiation, Synaptic Signalling and Neuronal Death (7 out of 20 significant GO terms) (Fig. 7b and Supplementary Table 3). Sashimi plots are shown for selected neuronal genes *Dynamin 1* (*Dnm1*), *Phosphatidylinositol Binding Clathrin Assembly Protein* (*Picalm*), *Protein Phosphatase 1 Regulatory Subunit 9A* (*Ppp1r9a*), *EFR3 homolog A* (*Efr3a*), *Very Low Density Lipoprotein Receptor* (*Vldlr*) and *Microtubule-Actin Crosslinking Factor 1* (*Macf1*) (Fig. 7c). Interestingly, *Picalm*, *Vldlr* and *Macf1* amongst others, have been previously identified as alternatively spliced upon *Tardbp* or *Fus* knowk-down in the mouse brain^28,29^. In addition, mis-splicing of *Picalm* has been reported in AD patients^30^, mutation-associated alternative splicing of *Dnm1* has been associated with epilepsy^31^ and mutations of *Efr3a* have been linked to autism spectrum disorder^32,33^. Taken together, the data indicate that *Sod1^G86R^* CSN undergo degeneration in a context of global RNA metabolism dysregulation and selective RNA splicing alterations that affect more particularly genes involved in proper neuronal development and function. These altered molecular pathways are reminiscent of other ALS-related genes which directly control RNA metabolism and splicing, *i.e. FUS* and *TARDBP* and point to common mechanisms across causative genes and neuronal subtypes, in, and above ALS context.

**Figure 7:**
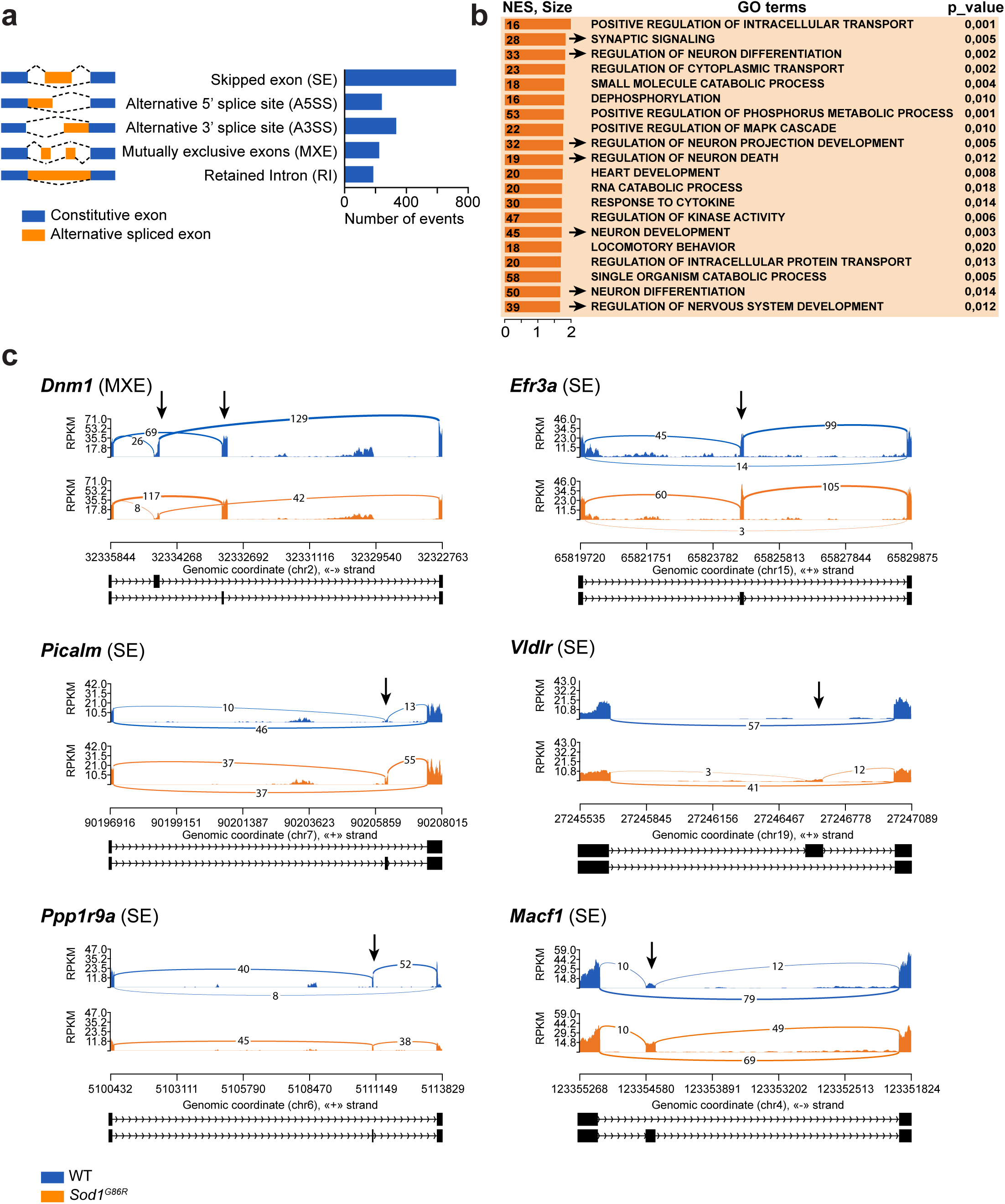
*Sod1^G86R^* CSN degeneration is correlated with numerous alternative splicing events that mostly affect neuronal genes. **a.** Schematic representation of the different alternative splicing events (left panel) identified between 90 day-old wild-type and *Sod1^G86R^* CSN samples, and the corresponding number of events for each type (right bar graph). **b.** Gene set enrichment analysis (GSEA) of 1,163 ranked genes for GO terms of Biological Processes. The 20 most enriched categories are listed according to their normalized enrichment score (NES) and p-value, and the numbers of corresponding genes (Size) are indicated. Arrows indicate the redundancy and overrepresentation of GO terms related to neuronal development and function. **c.** Sashimi plots representing alternative splicing events identified between wild-type (blue) and *Sod1^G86R^*(orange) CSN samples for four selected genes: *Dnm1*, *Picalm*, *Ppp1r9a*, *Efr3a*, *Vldlr* and *Macf1*. Chromosomal positions are indicated below each graph, and relevant exons are represented as black boxes. Arrows indicate significant alternative splicing event.

## Discussion

In clinics, ALS diagnosis relies on evidences for combined degeneration of CSN and MN. While the origin and propagation of the disease between the two neuronal populations has always been debated^34^, number of studies reported on the somatotopic relationship between cortical and spinal impairments^35-37^, ruling out the possibility that CSN and MN degenerations are independent, and rather supporting a propagation from one neuronal population to the other, whether descendent, from the cerebral cortex to the spinal cord, or ascendant, from the spinal cord to the cerebral cortex (not to be confused with the Wallerian or retrograde degeneration of the MN, from their axonal terminals to their cell bodies). In spite of its classical text book definition, ALS is highly heterogeneous and, amongst other parameters, relative upper (CSN) and lower (MN) motor neuron involvement and site of disease onset greatly vary across ALS patients^38^. Notably, *SOD1* mutant carriers typically present with spinal onset, which for the vast majority of them corresponds to a lower limb onset, and mild upper motor neuron impairment^38,39^. Similarly, mutant *SOD1* mouse models typically present with initial hind limb impairment, and so do the *Sod1^G86R^* mice^18,19^. Here, we took advantage of this well characterized model of the disease to test whether mice could recapitulate the somatotopic relationship between CSN and MN, and contribute to unravel the origin of the degeneration.

Because hyperreflexia and hypertonia that characterize CSN degeneration in Human^5,6^ are extremely difficult to assess in rodents, histology has become the standard to evaluate the CSN population in ALS mouse models^12-17^. Here, we chose retrograde labelling approaches to precisely label either the whole CSN population (Fig. 1, 2 and Supplementary Fig. 1), or the lumbar-projecting CSN subpopulation (Fig. 3 and Supplementary Fig. 3), as opposed to IHC, ISH or genetic approaches that allow for labelling of all cortical layer V subcerebral projection neurons amongst which CSN represent only a minority. When combined with rigorous sampling and counting, retrograde labelling further enabled to test whether sub-populations of CSN, that in rodent are topographically organized along both the medio-lateral and the rostro-caudal axes^21,22^, may be more affected by the disease. We show here that *Sod1^G86R^* mice displayed progressive loss of the whole CSN population (Fig.1 and Supplementary Fig. 2), and that, along the rostro-caudal axis, CSN loss started caudally and progressed rostrally (Fig. 2). Caudally-located CSN mostly project to the lumbar spinal cordand control hind limb movements^21,22^. Quantification of the lumbar-projecting CSN in *Sod1^G86R^* mice during the course of the disease indicated a significant loss as early as 60 days of age (Fig. 3), 45 days prior to MN cell bodies shrinkage and loss (Fig. 3 and Supplementary Fig. 4), 30 days prior to NMJ denervation, and 15 days prior to onset of weight loss, the first clinical symptom reported in this mouse line^40^. Together, the data indicate that, similarly to what has been reported for ALS patients, CSN and MN degeneration occur in a somatotopic manner in *Sod1^G86R^* mice, with CSN and MN involved in hind limb motor function being affected prior to the other subpopulations. In addition, CSN loss occurs before the first signs of MN degeneration (NMJ denervation), suggesting that, in this model, degeneration likely follows a descending propagation, from the motor cortex to the spinal cord. The data support the corticofugal hypothesis which arose from post-mortem histopathological analyses of brains from ALS brains of patients, and alive DTI studies^8-10^. Interestingly, in spite of the early onset of CSN loss, as many as 66.7% of the whole CSN population and 62.9% of the lumbar-projecting population remain at disease end-stage, indicating the existence of both disease-resistant and disease-vulnerable CSN. In the context of the corticofugal hypothesis, an important question to address in future studies would be to determine which of the loss of the most vulnerable CSN or the maintenance of disease-resistance CSN is the most detrimental to downstream MN, in other words, whether disease propagation from the cerebral cortex to the spinal cord arises from the loss of beneficial input or rather the maintenance of toxic input.

Transcriptomics, amongst other techniques, has largely contributed to our understanding of the molecular pathways leading to MN degeneration^41,42^. Similarly, we reasoned that transcriptomic analysis of CSN purified from adult *Sod1^G86R^* and control mice would unravel the molecular signalling that accompanies CSN degeneration in ALS. To this end, we adapted a method of purification of embryonic and young postnatal cortical neurons^20,25^ to adult cortical neurons under both healthy and neurodegenerative conditions, based on FAC Sorting of retrogradelly labelled neurons (Fig. 4). We thus purified CSN and CPN from the cerebral cortex of wild type and *Sod1^G86R^* mice at four disease-relevant adult ages, and conducted RNAseq analysis. To our knowledge, this is the first report on whole projection neuron population purification by FACS from the adult mouse cerebral cortex. In addition, the adapted methodology offers the advantage to utilise the neurons of individual mouse brains as true replicates, instead of pooling the neurons from several brains^20,43^, providing an adapted response to both ethical and statistical requirements. CPN isolated from same mouse brains were chosen as control, for their high similarity with CSN, and their resistance to degeneration (Supplementary Fig. 5) in spite of similar levels of *Sod1^G86R^* transgene expression (Fig. 5b). Possible future applications of this current method may prove useful to compare gene signatures of affected CSN populations amongst various ALS mouse models, or to highlight the differences between ALS and primary lateral sclerosis (PLS) characterized by CSN degeneration without MN impairment, potentially shedding light on putative propagation mechanisms from CSN to MN in ALS. Beyond this first attempt to unravel CSN alterations in ALS at the level of the population, single cell approaches will be, in the future, instrumental in identifying the mechanisms that discriminate disease-vulnerable from disease-resistant CSN, and contribute to inform the development of therapeutic strategies aimed at maintaining pools of properly functioning CSN^44^.

Analysis of the transcriptomic data first confirmed the selective degeneration of the CSN population and the minor impact of the *Sod1^G86R^* transgene expression on the CPN population (Fig. 5), in accordance with the histological quantifications (Fig. 1, 3, and Supplementary Fig. 2 and 5). Comprehensive analysis of gene expression data highlights pathways characteristic of ALS^1,2^, such as altered mitochondrial functions, protein folding and catabolism (Fig. 6a). Interestingly, the over-representation of RNA metabolism, including RNA splicing, amongst the most deregulated biological processes, cellular components and molecular functions, further confirmed by high throughput qPCR (Fig. 6) and splicing analysis (Fig. 7) is altogether expected in the context of ALS^1,2^, and novel for a *Sod1^G86R^* mutation. Indeed, altered RNA metabolism is reminiscent of other ALS-related genes which encode RNA binding proteins, such as *TARDBP* and *FUS* and whose knock-downs in the mouse brain are associated with altered RNA splicing^28,29^. Importantly, alternative splicing is also a hallmark of *C9ORF72*-associated mutations and of sporadic ALS^45^, and up-regulation of genes involved in mRNA processing and in RNA metabolic processes were also described in the cerebral cortex of sporadic ALS patients^46^. Gene set enrichment analysis of the statistically significant altered splicing events between WT and *Sod1^G86R^* CSN unravelled a large representation of genes involved in neuronal development and differentiation, synaptic signalling and neuronal death (Fig. 7b). Again, some of the targeted genes are reminiscent of those affected by *TARDBP* and *FUS* mutations^28,29^, such as the representative *Picalm*, *Vldlr* and *Macf1* (Fig. 7), highlighting altered neuron-specific functions (Supplementary Table 3). The mis-splicing of these transcripts, amongst others, could contribute to altered dendritic shape, synaptic transmission and intrinsic excitability of the CSN/layer V subcerebral projection neurons, which have been reported in various mouse models of the disease^14,16,17,47^, and potentially to overall motor cortex hyperexcitability, an early feature of cortical impairment in ALS patients^11^. Thus, beyond the *Sod1^G86R^* mouse model, the data are relevant to ALS in its broad genetic heterogeneity. This very first molecular temporal sequence of degenerating CSN paves the way to the identification of new therapeutic targets to prevent CSN loss, maintain their proper functioning, and potentially hinder disease spreading to downstream MN.

## Online Methods

### Animals

All animal experiments were performed under the supervision of authorized investigators and approved by the local ethical committee of Strasbourg University (CREMEAS, agreements # 00766.01 and 1534-2015082617403748). BAC transgenic male mice with the G86R murine *Sod1* missense mutation^18^ were obtained from the animal facility of the Faculty of Medicine, University of Strasbourg. Non-transgenic age-matched male littermates served as controls. Mice received water and regular rodent chow *ad libitum*. *Sod1^G86R^* animals were followed daily, and disease progression was rated according to a clinical scale going from score 4 to 0, as we previously described^19^. End-stage animals were harvested upon reaching score 0, *i.e.* when they were no longer unable to roll over within 10 s after being gently placed on their back^19^.

### Retrograde labelling of CSN

Mice were deeply anesthetized with intraperitoneal (i.p.) injection of Ketamine (Imalgène 1000®, Merial; 120 mg/kg body weight) and Xylazine (Rompun 2%®, Bayer; 16 mg/kg body weight) solution and placed on a heating pad. A precise laminectomy was performed in cervical (C3-C4) or lumbar (L1-L2) region of the spinal cord, depending on the experiment, and the animals were positioned below an injector (Nanoject II, Drummond Scientific, PA) mounted on a micromanipulator. Using pulled glass capillaries, the dura was punctured, the capillary lowered to the dorsal funiculus and five pressure microinjections of 23 nl of Fluorogold (Fluorchrome, CO) were performed on each side of the dorsal funiculus. Five days after Fluorogold injection, or upon reaching end stage of the disease, the animals were transcardially perfused with cold 0.01 M PBS, followed by cold 4% PFA in 0.01 M PBS and their brains, spinal cords, tibialis anterior and gastrocnemius muscles were collected.

### CSN quantification

Brains from Fluorogold injected animals were cut in 40 µm thick coronal sections on a Leica VT 1000S vibratome (Leika Biosystems) and mounted on slides with DPX mounting solution (SIGMA, 06522-100ML). Sections were imaged under an AxioImager.M2 fluorescence microscope (Zeiss) equipped with a structured illumination system (Apotome, Zeiss) and a highly sensitive black and white camera (Hamamatsu). Images were acquired using the ZEN 2 software (Zeiss). For mice injected in the cervical portion of the spinal cord, all FG-positive neurons present on both hemispheres of 8 equally spaced coronal sections, separated by 480 µm and spanning from 2.10 to −1.70 relative to Bregma, were counted (Supplementary Fig. 1). For mice injected in the lumbar portion of the spinal cord, all FG-positive neurons present on both hemispheres of 3 equally spaced coronal sections, separated by 480 µm and spanning from −0.46 to −1.70 relative to Bregma, were counted (Supplementary Fig. 2). CSN quantification was performed using a macro that we developed on ImageJ (1.48k, NIH) by two experimenters blinded to the age and genotype of the samples. In brief, this macro generates a reversed image and performs a noise reduction with the “Median filter” (radius=2). A Gaussian blur (sigma=25) filtered image is then created and subtracted from the non-blurred image. The result is handled by the “rolling ball radius Subtract Background” algorithm (rolling=18). The edges of the cells are sharpened and enhanced by the “Unsharp Mask filter” (radius=50 mask=0.6). The user then can adjust the automatically set threshold that will be used for the determination of black and white pixels of the binarized image version. The image is then binarized with the “Make Binary” function and subjected to several morphological operations to close (“Close-“) and fill holes (“Fill Holes”) in cells and eliminate small objects (“Erode” and “Dilate”) and separate touching cells by “Watershed” segmentation. If needed, the user can further separate touching cells (white line drawing between cells), complete the filling of the holes (black brush drawing) and define a ROI around the whole cell population to avoid image border artefacts to be counted. Cells are then counted with the “Analyse particles” function. The macro automatically exports the results as a spreadsheet file and an image file.

The cDNA clone for *Crystallin-mu* (Crym) is a kind gift from the Arlotta lab. Riboprobes were generated as previously described^20^. Nonradioactive *in situ* hybridization was performed on 40 µm vibratome coronal brain sections 0.14 mm anterior to Bregma. Selected sections were mounted on superfrost slides and processed using reported methods^48^. Bright field 10X tiles images were acquired with a Nikon microscope equipped with a Nikon Digital Sight DS-U3 camera and run by the Nis Elements 4.0 software. *Crym*-positive neurons were manually counted, on both hemispheres, within layer V, from M2 medially to S1 laterally by two experimenters blinded to the age and genotype of the samples.

### Spinal motor neurons quantification

The L1-L4 lumbar level of the spinal cords were cut on a vibratome into coronal sections of 40 µm. 4 nonadjacent sections spaced by 320 µm were labelled by immunohistochemistry as previously described^49^, using a goat anti-choline acetyltransferase (ChAT) antibody (Millipore) and a biotinylated donkey anti-goat IgG (Jackson). Two images per section (one per ventral horn) were captured using an AxioImager.M2 microscope (Zeiss) equipped with a high-resolution B/W camera (Hamamatsu) and run by the ZEN 2 software (Zeiss). Cells bodies sizes were measured using ImageJ (NIH).

### Electromyography

Mice were anesthetized with a solution of Ketamine/Xylazine (100 mg/kg; 5 mg/kg) and kept under a heating mat to maintain a physiological muscle temperature (±31°C). Electrical activity was monitored on the muscles of both limbs for at least 2 min as previously described^19^.

### Neuromuscular junctions staining and morphological analysis

Tibialis anterior muscles were dissected into bundles and processed for immunofluorescence with a rabbit anti-synaptophysin antibody, a rabbit anti-neurofilament antibody (Eurogentec) followed by Alexa-conjugated donkey anti-rabbit 488 (Jackson) and rhodamine-conjugated α-bungarotoxin (Sigma-Aldrich), as previously described^49^. Neuromuscular junctions (NMJs) analysis was performed by an independent age and genotype-blinded observer, directly under and AxioImager.M2 microscope (Zeiss) equipped with a high-resolution B/W camera (Hamamatsu) and run by the ZEN 2 software (Zeiss). On average 500 NMJs per animals were examined. NMJs were considered partially denervated when the presynaptic nerve terminal labelled with synaptophysin was partially observed from the postsynaptic region labelled with α-bungarotoxin, and denervated when the presynaptic nerve terminal was totally absent. Representative NMJs were imaged using the same microscope setting.

### Statistical analyses

Data are expressed as the means ± SEM (standard error of the mean). Statistical analyses were performed using GraphPad Prism 6 (GraphPad, CA). Multiple t tests corrected for multiple comparisons using the Holm-Šídák method were used to analyse CSN and MN quantifications and occurrences of denervated NMJ and positive EMG recordings.

### CSN and CPN purification

For FACS experiments, CSN from *Sod1^G86R^* mice and their wild-type littermates were retrogradelly labelled using Green IX Retrobeads (Lumafluor) injected at 25 days of age with the same protocol as for Fluorogold injection. Purification of CSN and CPN were performed in parallel from same WT or *Sod1^G86R^* mice at 30, 60, 90 and 105 days of age. Mice were deeply anesthetized with by an i.p. injection of Ketamine/Xylazine (100 mg/kg; 5 mg/kg) before decapitation. The brains were sectioned in a stainless steel coronal brain matrix (Harvard Apparatus, MA), and 1 mm-thick sections were transferred under a fluorescence SMZ18 microscope (Nikon). The layer V containing labelled CSN and the upper layer II/III above were microdissected from 4 coronal sections and collected in separate tubes filled with iced HABG (Hibernate A (BrainBits UK), B27, Glutamax (Gibco) and 0.1N NaOH). The microdissected tissues were enzymatically digested with 34 U/ml papain at 37°C for 30 min. Cells were mechanically dissociated by gentle trituration in iced HABG, filtered through 70 μm cell strainer (BD Falcon) and subjected to density centrifugation through a three-density step gradient of Percoll (Sigma, MO) as previously described^50^. Upon centrifugation, the cell pellets were resuspended in cold 0.01 PBS and fixed with 70% EtOH for 30 min at 4°C. Fixed cells were centrifugated to eliminate EtOH and resuspended in 0.01 M PBS complemented with RNAse inhibitors (Promega). Microsphere-labelled CSN were purified using the FACS AriaII (BD Biosciences), based on their fluorescence, size and granularity. Unlabelled CPN were purified by FACS, based on their size and granularity. Approximately 2 000 CSN and exactly 2 000 CPN were be obtained from each adult mouse brain and were used as individual biological replicates for RNA sequencing. Upon purification, neurons were centrifugated and frozen as cell pellets.

### RNA sequencing

Full length cDNA were generated from frozen cell pellets using the SMART-Seq v4 Ultra Low Input RNA kit for Sequencing (Clontech, CA) according to the manufacturer’s instructions. 11 cycles of cDNA amplification were performed using the Seq-Amp polymerase. 600 ng of pre-amplified cDNA were then used as input for Tn5 transposon tagmentation by the Nextera XT kit (Illumina) followed by 12 cycles of library amplification. Following purification using Agencourt AMPure XP beads (Beckman Coulter), the size and concentration of the libraries were assessed on an Agilent 2100 Bioanalyzer. The libraries were then loaded in the flow cell at a concentration of 3 nM, clusters were generated by using the Cbot and sequenced on the Illumina HiSeq 4000 system as paired-end 2×50-base reads, following Illumina’s instructions.

### Data repository

RNAseq data have been deposited in the ArrayExpress database at EMBL-EBI under accession number E-MTAB-7876 (https://www.ebi.ac.uk/arrayexpress/experiments/E-MTAB-7876/).

### RNA sequencing, gene set enrichment, gene network and differential alternative splicing analyses

Reads were mapped onto the mm10 assembly of mouse genome using Tophat v2.0.14^51^ and bowtie v2.1.0 aligner. Gene expression was quantified using HTSeq v0.6.1^52^ and gene annotations from Ensembl release 84. Hierarchical clustering of samples was calculated using the UPGMA algorithm (Unweighted Pair Group Method with Arithmetic Mean) with the SERE coefficient as the distance measure^53^. Comparisons between the *Sod1^G86R^* and WT CSN, and between *Sod1^G86R^* and WT CPN samples across ages, and between *Sod1^G86R^* and WT CSN at each age were performed using the method proposed by Love et al.^54^ implemented in the DESeq2 Bioconductor library (DESeq2 v1.6.3). Thresholds of |log2FoldChange| > 1 and adjusted p value < 0.05 were chosen to identify significantly differentially expressed genes. GSEA^26^ was performed on a pre-ranked list of 29,017 genes significantly dysregulated in CSN and not CPN, and with an FPKM >14 in more than 90% of samples, using −log_10_ of the pvalue x sign of the log_2_Fold-Change as ranking metric, and the gene set database of Biological Process Ontology (version 2.5) from the MSigDB collections. Enrichment Maps were generated with Cytoscape^55^. Differential alternative splicing analysis was performed using rMATS version 3.2.5^27^ on alignments performed with STAR 2.5.3a^56^, and splicing with reads that span splicing junctions were considered for further analysis, using the threshold of FDR < 0.05. GSEA analysis was performed on a list of 1,163 mis-spliced genes, using −log10 of the FDR as ranking metric. Sashimi plots were generated with rmats2sashimiplot^27^.

### Fluidigm Validation

High-throughput qPCR validation was carried out on the Fluidigm’s Biomark HD system, according to the manufacturer’s protocol. Specifically, 100 ng of RNA were reverse-transcribed using the Reverse Transcription Master Mix Kit (Fluidigm). cDNA was then pre-amplified using 50 nM pooled primer mixture and the PreAmp Master Mix Kit (Fluidigm) for 14 cycles. Unincorporated primers were removed with 8U of Exonuclease I (NEB) and the reaction products were diluted 5-fold in TE Buffer. The treated cDNA’s were combined with SsoFast EvaGreen Supermix with Low ROX (Bio-Rad) and loaded onto a 96.96 Dynamic Array integrated fluidic circuit (Fluidigm). Primers used were:

*Aars*, F: TGGTTTCTGGTGGACTTGCAG - R: AAGTCTCCTCGGGAACCTTAGC

*Dars*, F: TCGAGTTAAGGACCTGACAGT - R: CACTGCTTCCCTTTTGCTCT

*Ddx20*, F: TGCACAGCAGAGCTCAACAT - R: GCATCAAGGCGCTGATTCTG

*Eif2b5*, F: GCCACCAACAGGGTTCTTCA - R: CCTGGAACAGGCTCAATGGAA

*Eif3h*, F: AGATCGACGGCCTGGTAGTAT - R: CCTGAACAACCTCAGTGCCT

*Eif4a2*, F: TAGTATTGGCCCCCACCAGA - R: AGCATGACAAGTTGCTCCCA

*Eif4g2*, F: TTATGAAAAGCCAGGGGCTAAG - R: CCGAGGTGGCATATCCTTCG

*Gemin8*, F: CAACTGGAGGGCTAGTGCAT - R: GCATCCAAAGCATTGCGTGAT

*Kars*, F: AGCAAGAAGTGGTGGTAATCAGT - R: CCCTAACAAGCCTGACAGCA

*Nsun2*, F: GAGCGATGCCTTCGAATCCT – R: GACGTTTGTTCCACGGCATT

*Ppprf31*, F: CAGCTCGTGTGGACAGCTT - R: GCTCAATCTCATCCTTTAGTTCGT

*Sars*, F: CGTTTCTCTACACAGAACAAGTTGA - R: GGACAAACTCCACCTTCTTGG

*Sf3bp4*, F: GGTGCTGCTGCTTAGAGACG – R: TACACCGTGGCATCCTGATT

*Slbp*, F: CCTGAAAGCTTTACAACTCCTGA – R: ATCTTCTTCAACTGCACTGGC

*Snrpb*, F: CCCACCTCCAGGCATTATGG - R: TGCCTATAGGTGTCCCACGA

*Trnt1*, F: GTAAGCGGGTCTTCGGAGTA – R: CGCCAATCACACATGCTGC

## Acknowledgements

The work has been supported by an ERC starting grant #639737, a Marie Curie career integration grant #618764, an AFM-Telethon trampoline grant #16923 and a Neurex grant to CR, as well as an Inserm/Région Alsace PhD fellowship to CM. Sequencing was performed by the GenomEast platform, a member of the “France Génomique” consortium (ANR-10-INBS-0009). The authors are thankful to Drs Luc Dupuis, Jose-Luis Gonzalez de Aguilar, Simona Lodato and Guillaume Tena for critical reading of the manuscript and insightful comments.

---

**Supplementary Figure 1: Retrograde labelling of CSN upon Fluorogold injection in the cervical portion of the spinal cord**

Representative negative fluorescence images of brain coronal sections numbered 1 (Bregma 2.58 mm) to 10 (Bregma −2.30 mm) showing Fluorogold-labelled CSN in the cerebral cortex of a 90 day-old wild-type mouse. Upon Fluorogold injection into the cervical portion of the dorsal funiculus, labelled CSN were consistently observed from section 2 (Bregma 2.10 mm) to 9 (Bregma −1.70 mm). Scale bar: 1 mm.

**Supplementary Figure 2: The population of *Crym*-positive neurons progressively disappears from the layer V of the motor cortex of *Sod1^G86R^* mice**

**a.** Representative images of *in situ* hybridization on brain coronal sections (Bregma 0.14 mm) of end stage *Sod1^G86R^* and 115 day-old wild-type control mice showing decreased *Crym* expression in the layer V of the cerebral cortex. Dotted red rectangles indicate the areas where *Crym*-positive neurons were counted. **b.** Close-ups of the M1 area. **c.** Bar graph representing the percentage of *Crym*-positive neurons in *Sod1^G86R^* mice (orange) relative to their wild-type littermates (grey) at 60, 90, 105 and 115 days (d) of age or end stage (ES). * p<0.05; ** p<0.01. Scale Bar = 200µm in a. and 100µm in b.

**Supplementary Figure 3: Retrograde labelling of CSN upon Fluorogold injection in the lumbar portion of the spinal cord**

Representative negative fluorescence images of brain coronal sections numbered 1 (Bregma 2.58 mm) to 10 (Bregma −2.30 mm) showing Fluorogold-labelled CSN in the cerebral cortex of a 90 day-old wild-type mouse. Upon Fluorogold injection into the lumbar portion of the dorsal funiculus, labelled CSN were consistently observed from section 7 (Bregma −0.46 mm) to 9 (Bregma −1.70 mm). Scale bar: 1 mm.

**Supplementary Figure 4: In *Sod1^G86R^* mice, NMJ denervation starts after 75 days**

**a.** Bar graph representing the proportion of animals with electromyography (EMG) recordings of denervated (orange) or innervated (grey) muscles. **b.** Representative electromyography recordings of innervated (left) and denervated (right) muscles. **c.** Bar graph representing the average proportions of innervated (grey), partly denervated (orange) of fully denervated (red) neuromuscular junctions (NMJ) upon staining and microscopy analysis of one tibialis anterior muscle of wild-type *Sod1^G86R^* mice at 60, 75, 90 and 105 days of age or end stage (ES). **d.** Representative maximum intensity projection images of z-stacks of typical innervated, partly or fully denervated NMJ from 90 day-old *Sod1^G86R^* and wild-type mice. * p<0.05; *** p<0.001 in multiple t-test. Scale Bar = 10 µm.

**Supplementary Figure 5: In *Sod1^G86R^* mice, the number of CUX-2 positive CPN remains stable over time**

**a.** Representative fluorescence images of CUX2 staining on brain coronal sections from 60 day-old wild-type and *Sod1^G86R^* mice. **b.** Representative close-ups of CUX2 staining on brain coronal sections from wild-type and *Sod1^G86R^* mice of 60, 75, 90 or 105 days (d) of age of end-stage (ES). **c.** Bar graph representing the average number of CUX2-positive nuclei on close-ups of matched brain coronal sections from *Sod1^G86R^* (orange) and wild-type mice (grey). *** p<0.001 in multiple t-tests. Scale bars: 200 µm in a. and 50 µm in c.

**Supplementary Figure 6: Enrichment map.**

Nodes represent gene sets, and their size is proportional to the number of corresponding genes. Edges represent overlaps between gene sets, and their width is proportional to the number of overlapping genes. Node colour and intensity vary with the corresponding NES: orange for positive and blue for negative NES respectively, the darker the higher. The dotted line circles the GO terms linked to RNA metabolism.

**Supplementary Table 1**

GSEA analysis of transcriptomic data: Gene Ontology (GO) terms for Biological Processes (BP), Cellular Components (CC) and Molecular Functions (MF).

**Supplementary Table 2**

Genes involved in RNA metabolism and already linked to motoneuron and/or neurodevelopmental diseases.

**Supplementary Table 3**

GSEA analysis of 1,163 ranked mis-spliced genes, for the GO terms Biological Process, Cellular Component and Molecular Function.

## References

1. Brown, R. H. & Al-Chalabi, A. Amyotrophic Lateral Sclerosis. N. Engl. J. Med. 377, 162–172 (2017).

2. van Es, M. A. et al. Amyotrophic lateral sclerosis. The Lancet 390, 2084–2098 (2017).

3. Taylor, J. P., Brown, R. H. & Cleveland, D. W. Decoding ALS: from genes to mechanism. Nature 539, 197–206 (2016).

4. Weishaupt, J. H., Hyman, T. & Dikic, I. Common Molecular Pathways in Amyotrophic Lateral Sclerosis and FrontotemporalDementia. Trends Mol. Med. 22, 769–783 (2016).

5. Eisen, A. et al. Cortical influences drive amyotrophic lateral sclerosis. J. Neurol. Neurosurg. Psychiatry 88, jnnp–2017–315573 (2017).

6. Geevasinga, N., Menon, P., Özdinler, P. H., Kiernan, M. C. & Vucic, S. Pathophysiological and diagnosticimplications of cortical dysfunctionin ALS. Nat. Rev. Neurol. 1–11 (2016).

7. Ling, S.-C., Polymenidou, M. & Cleveland, D. W. Converging Mechanisms in ALS and FTD: Disrupted RNA and Protein Homeostasis. Neuron 79, 416–438 (2013).

8. Braak, H. et al. Amyotrophic lateral sclerosis-a model of corticofugal axonal spread. Nat. Rev. Neurol. 9, 708–714 (2013).

9. Brettschneider, J. et al. Stages of pTDP-43 pathology in amyotrophic lateral sclerosis. Ann Neurol. 74, 20–38 (2013).

10. Kassubek, J. et al. Diffusion tensor imaging analysis of sequential spreading of disease in amyotrophic lateral sclerosis confirms patterns of TDP-43 pathology. Brain 137, 1733–1740 (2014).

11. Vucic, S. & Kiernan, M. C. Transcranial Magnetic Stimulationfor the Assessment of Neurodegenerative Disease. Neurotherapeutics 1–16 (2017).

12. Zang, D. W. & Cheema, S. S. Degeneration of corticospinal and bulbospinal systems in the superoxide dismutase 1. Neurosci. Lett. 332, 99–102 (2002).

13. Ozdinler, P. H. et al. Corticospinal Motor Neurons and Related Subcerebral Projection Neurons Undergo Early and Specific Neurodegeneration in hSOD1G93A Transgenic ALS Mice. J. Neurosci. 31, 4166–4177 (2011).

14. Yasvoina, M. V. et al. eGFP Expression under UCHL1 Promoter Genetically Labels Corticospinal Motor Neurons and a Subpopulation of Degeneration-Resistant Spinal Motor Neurons in an ALS Mouse Model. J. Neurosci. 33, 7890–7904 (2013).

15. Handley, E. E. et al. Synapse Dysfunction of Layer V Pyramidal Neurons Precedes Neurodegeneration in a Mouse Model of TDP-43 Proteinopathies. Cerebral Cortex 27, 3630–3647 (2017).

16. Fil, D. et al. Mutant Profilin1 transgenic mice recapitulate cardinal features of motor neuron disease. Hum. Molecul. Genet. 364, ddw429 (2016).

17. Liu, Y. et al. C9orf72 BAC Mouse Model with Motor Deficits and Neurodegenerative Features of ALS/FTD. Neuron 90, 521–534 (2016).

18. Ripps, M. E., Huntley, G. W., Hof, P. R., Morrison, J. H. & Gordon, J. W. Transgenic mice expressing an altered murine superoxide dismutase gene provide an animal model of amyotrophic lateral sclerosis. PNAS 92, 689–693 (1995).

19. Rouaux, C. et al. Sodium valproate exerts neuroprotective effects in vivo through CREB-binding protein-dependent mechanisms but does not improve survival in an amyotrophic lateral sclerosis mouse model. J. Neurosci. 27, 5535–5545 (2007).

20. Arlotta, P. et al. Neuronal subtype-specific genes that control corticospinal motor neuron development in vivo. Neuron 45, 207–221 (2005).

21. Ayling, O. G. S., Harrison, T. C., Boyd, J. D., Goroshkov, A. & Murphy, T. H. Automated light-based mapping of motor cortex by photoactivation of channelrhodopsin-2 transgenic mice. Nat. Methods 6, 219–224 (2009).

22. Kamiyama, T. et al. Corticospinal Tract Development and Spinal Cord Innervation Differ between Cervical and Lumbar Targets. J. Neurosci. 35, 1181–1191 (2015).

23. Fischer, L. R. et al. Amyotrophic lateral sclerosis is a distal axonopathy: evidence in mice and man. Exp. Neurol. 185, 232–240 (2004).

24. Molyneaux, B. J. et al. Novel subtype-specific genes identify distinct subpopulations of callosal projection neurons. J. Neurosci. 29, 12343–12354 (2009).

25. Ye, Z. et al. Instructing Perisomatic Inhibition by Direct Lineage Reprogramming of Neocortical Projection Neurons. Neuron 88, 475–483 (2015).

26. Subramanian, A. et al. Gene set enrichment analysis: A knowledge-basedapproach for interpreting genome-wideexpression profiles. PNAS 102, 15545–15550 (2005).

27. Shen, S., et al. rMATS: Robust and flexible detection of differential alternative splicing from replicate RNA-Seq data. PNAS 111, E5593–E5601 (2014).

28. Lagier-Tourenne, C. et al. Divergent roles of ALS-linked proteins FUS/TLS and TDP-43 intersect in processing long pre-mRNAs. Nat. Neuro. 15, 1488–1497 (2012).

29. Polymenidou, M. et al. Long pre-mRNA depletion and RNA missplicing contribute to neuronal vulnerability from loss of TDP-43. Nat. Neuro. 14, 459–468 (2011).

30. Raj, T. et al. Integrative transcriptome analyses of the aging brain implicate altered splicing in Alzheimer’s disease susceptibility. Nat. Genet. 50, 1584–1592 (2018).

31. Boumil, R. M. et al. A Missense Mutation in a Highly Conserved Alternate Exon of Dynamin-1 Causes Epilepsy in Fitful Mice. PLoS Genet. 6, e1001046 (2010).

32. Gupta, A. R. et al. Rare deleterious mutations of the gene EFR3A in autism spectrum disorders. 5, 1–14 (2014).

33. Marui, T. et al. Association of the neuronal cell adhesion molecule (NRCAM) gene variants with autism. Int. J. Neuropsychopharm. 12, 1 (2008).

34. Ravits, J. M. & La Spada, A. R. ALS motor phenotype heterogeneity,focality, and spread. Neurology 73, 805–811 (2009).

35. Mochizuki, Y., Mizutani, T., Shimizu, T. & Kawata, A. Neuroscience Letters. Neurosci. Lett. 503, 73–75 (2011).

36. Ravits, J., Paul, P. & Jorg, C. Focality of upper and lower motor neuron degeneration at the clinical onset of ALS. Neurology 68, 1571–1575 (2007).

37. Zhang, Q. et al. Side of Limb-Onset Predicts Laterality of Gray Matter Loss in Amyotrophic Lateral Sclerosis. Biomed Res. Int. 2014, 1–11 (2014).

38. Swinnen, B. & Robberecht, W. The phenotypic variability of amyotrophic lateral sclerosis. Nat. Rev. Neurol. 10, 661–670 (2014).

39. Millecamps, S. et al. SOD1, ANG, VAPB, TARDBP, and FUS mutations in familial amyotrophic lateral sclerosis: genotype-phenotype correlations. J. Med. Genet. 47, 554–560 (2010).

40. Dupuis, L., Oudart, H., Rene, F., Gonzalez de Aguilar, J.-L. & Loeffler, J.-P. Evidence for defective energy homeostasis inamyotrophic lateral sclerosis: Benefit of ahigh-energy diet in a transgenic mouse model. PNAS 1–6 (2004).

41. Caballero-Hernandez, D. et al. The ‘Omics’ of Amyotrophic Lateral Sclerosis. Trends Mol. Med. 1–15 (2015).

42. Ferraiuolo, L., Kirby, J., Grierson, A. J., Sendtner, M. & Shaw, P. J. Molecular pathways of motor neuron injury in amyotrophic lateral sclerosis. Nat. Rev. Neurol. 7, 616–630 (2011).

43. Molyneaux, B. J. et al. DeCoN: Genome-wide Analysis of In Vivo Transcriptional Dynamics during Pyramidal Neuron Fate Selection in Neocortex. Neuron 85, 275–288 (2014).

44. Kulkarni, A., Anderson, A. G., Merullo, D. P. & Konopka, G. ScienceDirectBeyond bulk: a review of single cell transcriptomics methodologies and applications. Cur. Opin. Biotechnol. 58, 129–136 (2019).

45. Prudencio, M. et al. Distinct brain transcriptome profiles in C9orf72-associated and sporadic ALS. Nat. Neurosci. 18, 1175–1182 (2015).

46. Andrés-Benito, P., Moreno, J., Aso, E., Povedano, M. & Ferrer, I. Amyotrophic lateral sclerosis, gene deregulation in the anterior horn of the spinal cord and frontal cortex area 8: implications in frontotemporal lobar degeneration. Aging 9, 823–851 (2017).

47. Kim, J. et al. Changes in the Excitability of Neocortical Neurons in a Mouse Model of Amyotrophic Lateral Sclerosis Are Not Specific to Corticospinal Neurons and Are Modulated by Advancing Disease. J. Neurosci. 37, 9037–9053 (2017).

48. Lodato, S. et al. Excitatory projection neuron subtypes control the distribution of local inhibitory interneurons in the cerebral cortex. Neuron 69, 763–779 (2011).

49. Scekic-Zahirovic, J. et al. Toxic gain of function from mutant FUS protein is crucial to trigger cell autonomous motor neuron loss. EMBO J. 35, 1077–1097 (2016).

50. Guez-Barber, D. et al. Journal of Neuroscience Methods. J. Neurosci. Methods 203, 10–18 (2012).

51. Kim, D. et al. TopHat2: accurate alignment of transcriptomes in the presence of insertions, deletions and gene fusions. Genome Biol 14, R36 (2013).

52. Anders, S., Pyl, P. T. & Huber, W. HTSeq — A Python framework to work with high-throughput sequencing data. Bioinformatics 31, 166–169 (2015).

53. Schulze, S. K., Kanwar, R., lzenleuchter, M. G., Therneau, T. M. & Beutler, A. S. SERE: Single-parameter quality control and sample comparison for RNA-Seq. BMC Genomics 13, 1–1 (2012).

54. Love, M. I., Huber, W. & Anders, S. Moderated estimation of fold change and dispersion for RNA-seq data with DESeq2. Genome Biol 15, 31 (2014).

55. Shannon, P. et al. Cytoscape: A Software Environment for Integrated Models of Biomolecular Interaction Networks. Genome e Reserach 13, 2498–2504 (2003).

56. Dobin, A. et al. STAR: ultrafast universal RNA-seq aligner. Bioinformatics 29, 15–21 (2012).

