## Supplementary Figures and Tables for "Early alterations of RNA metabolism and splicing from adult corticospinal neurons in an ALS mouse model"

Marques *et al.*, Supplementary Figure 1

#### FLUOROGOLD - Retrograde Labelling from the Cervical Spinal Cord

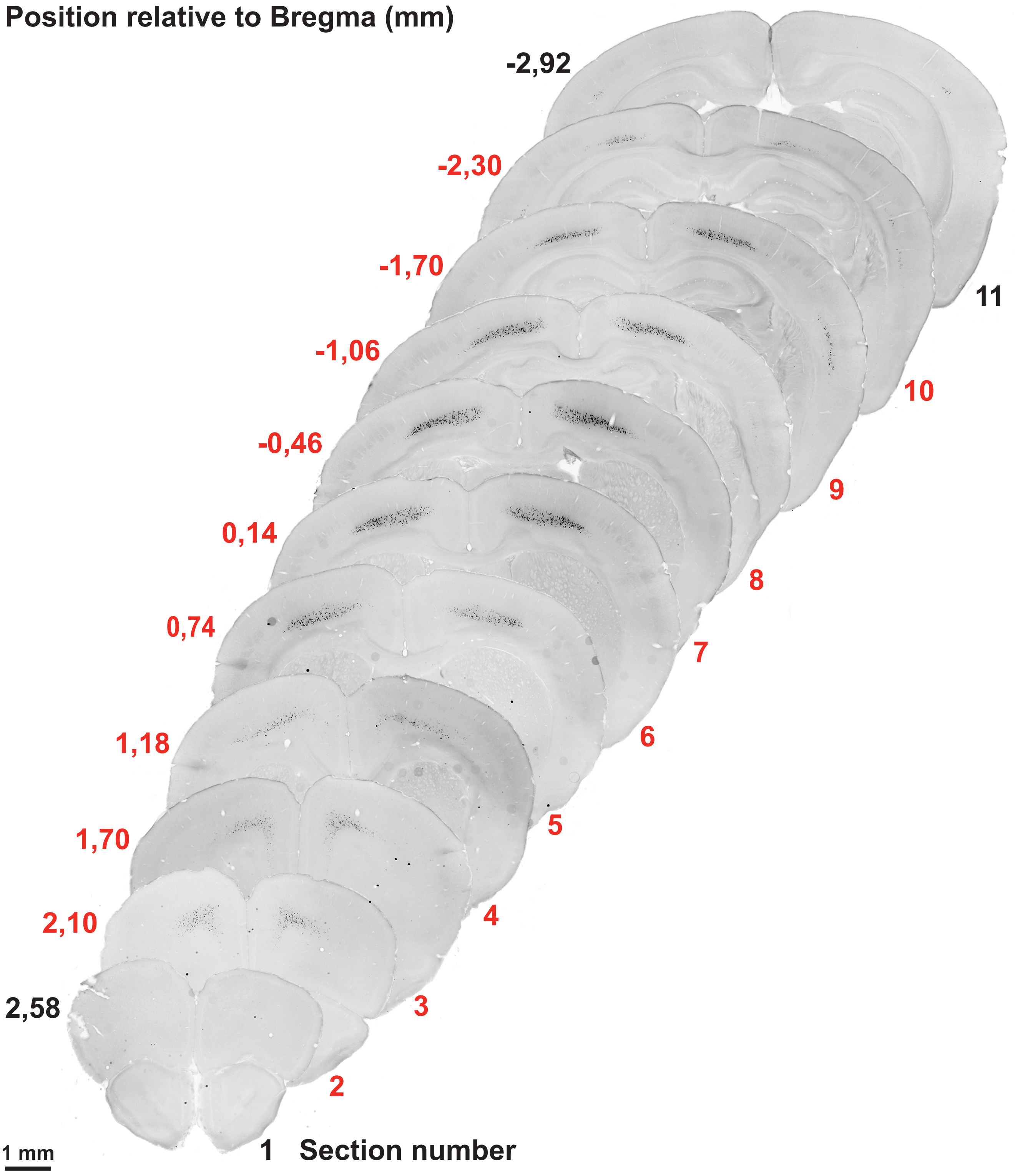

**Marques *et al.*, Supplementary Figure 2**

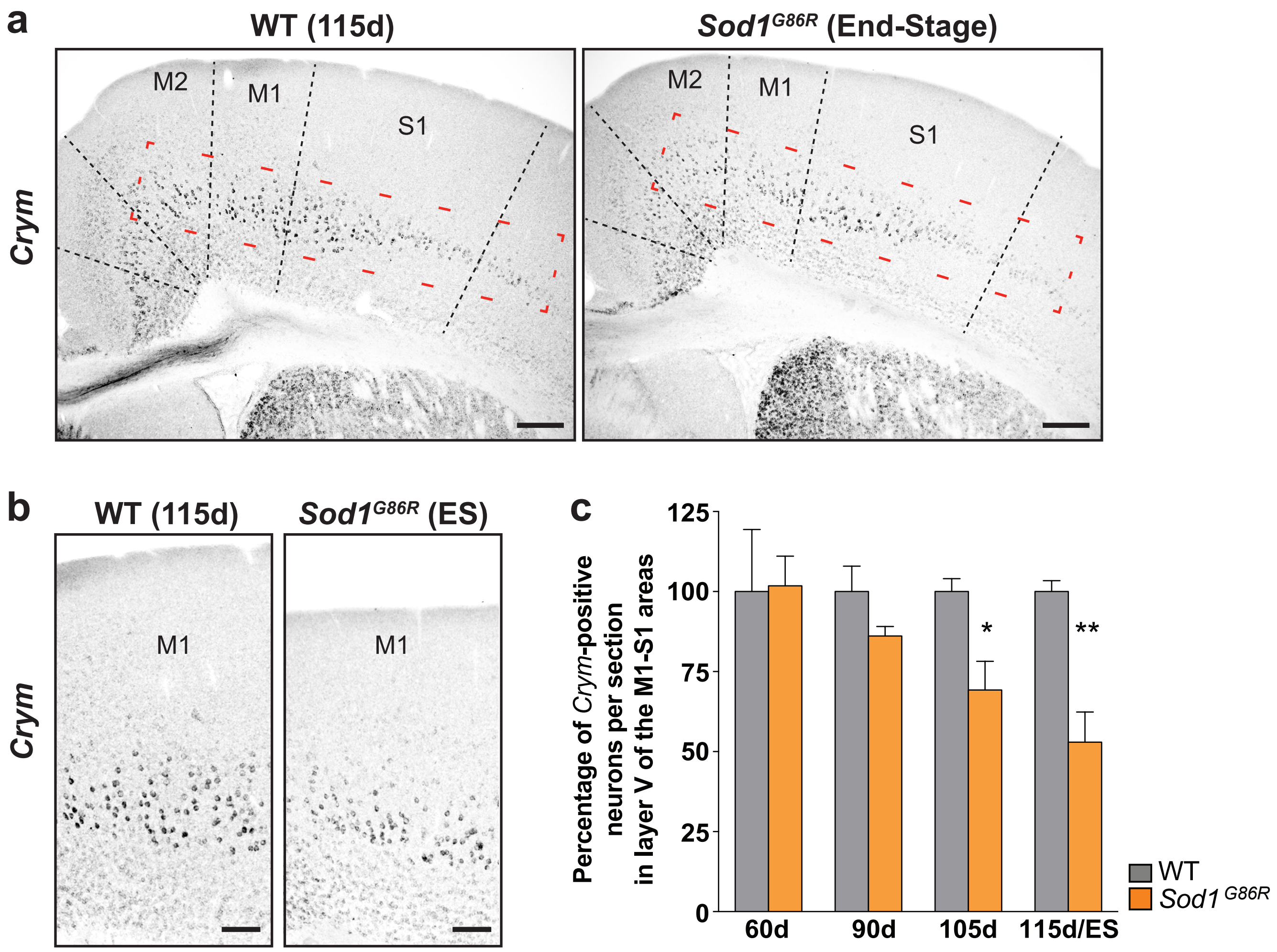

Marques *et al.*, Supplementary Figure 3

#### FLUOROGOLD - Retrograde Labelling from the Lumbar Spinal Cord

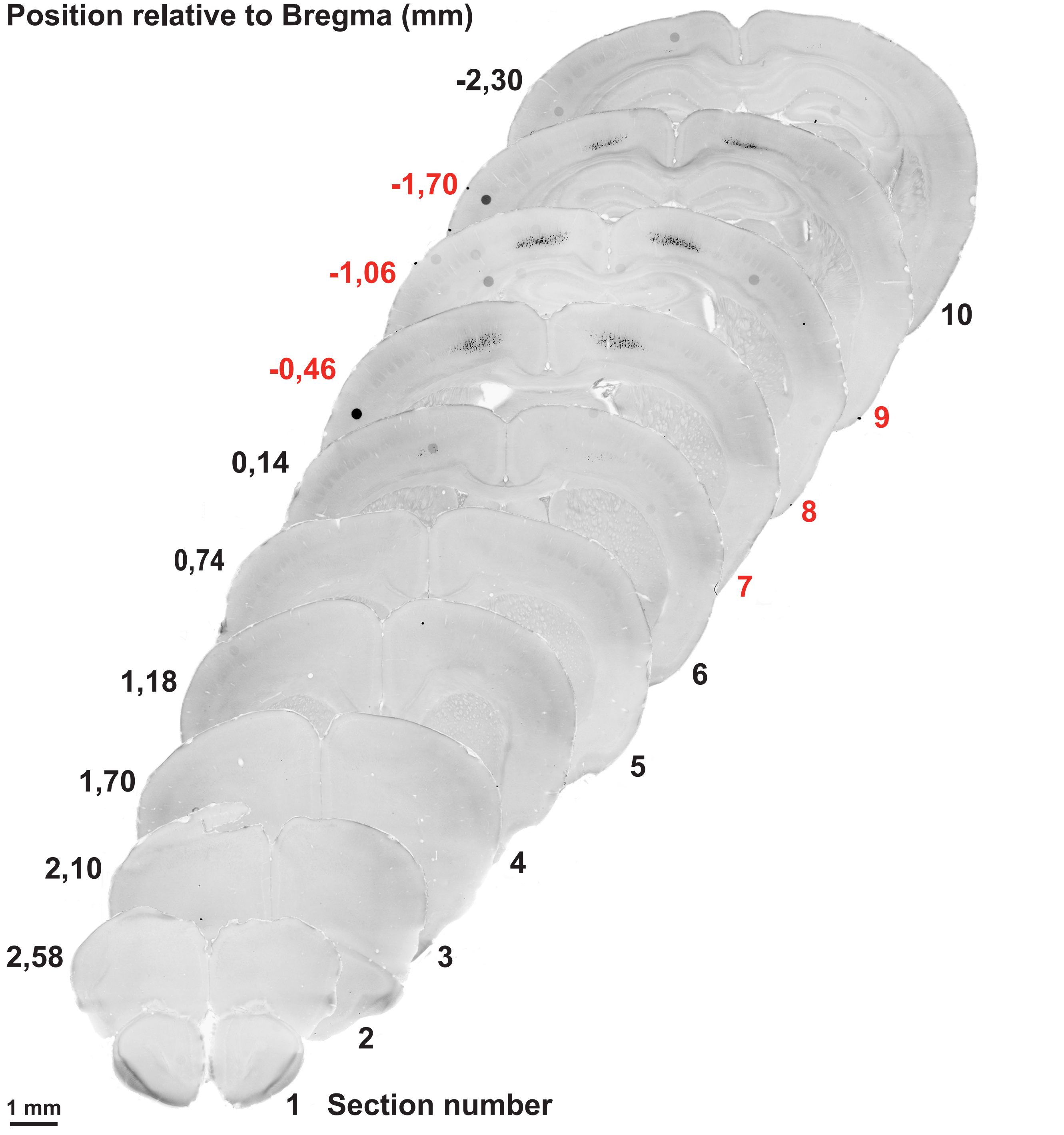

#### Marques *et al.*, Supplementary Figure 4

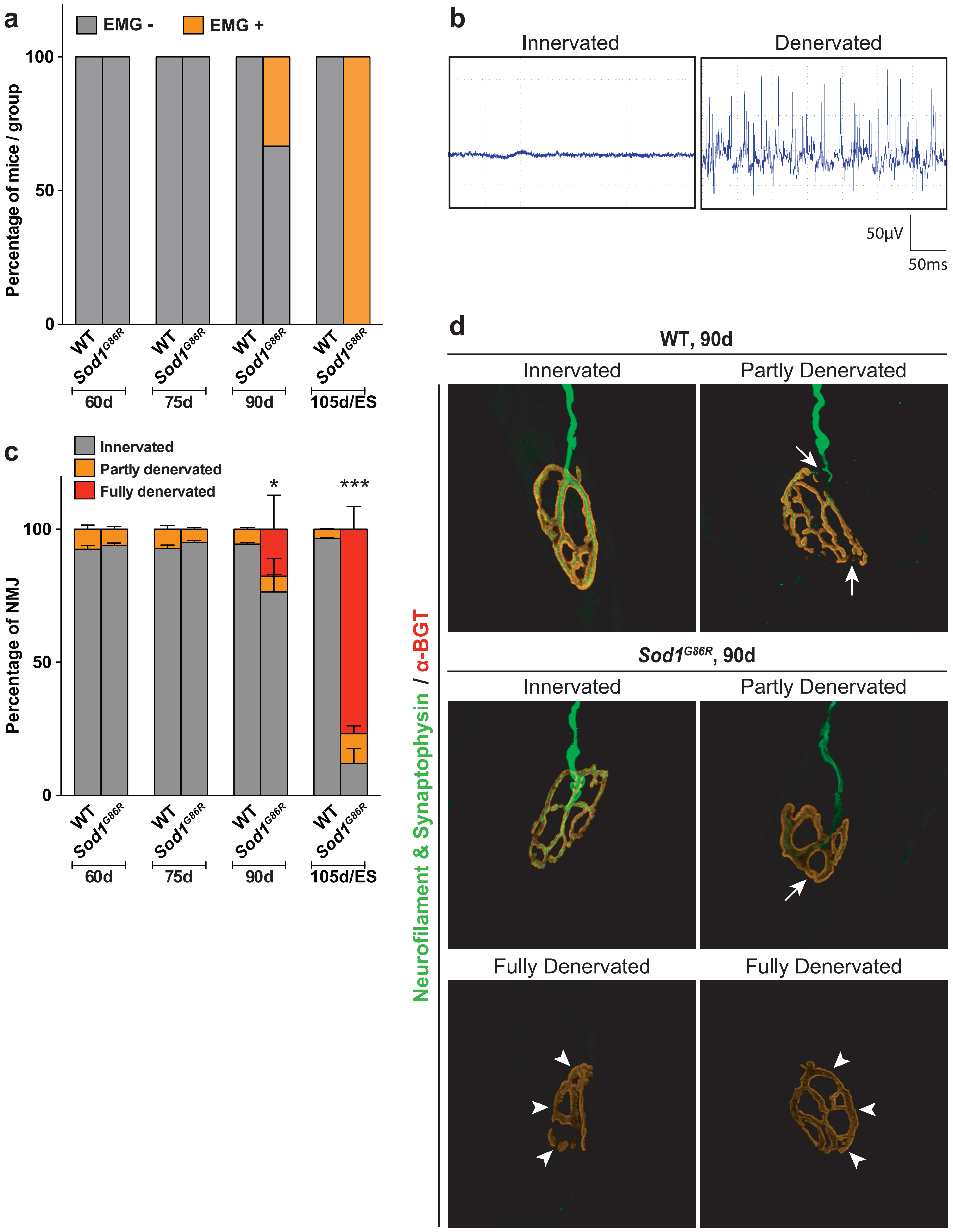

Marques *et al.*, **Supplementary Figure 5**

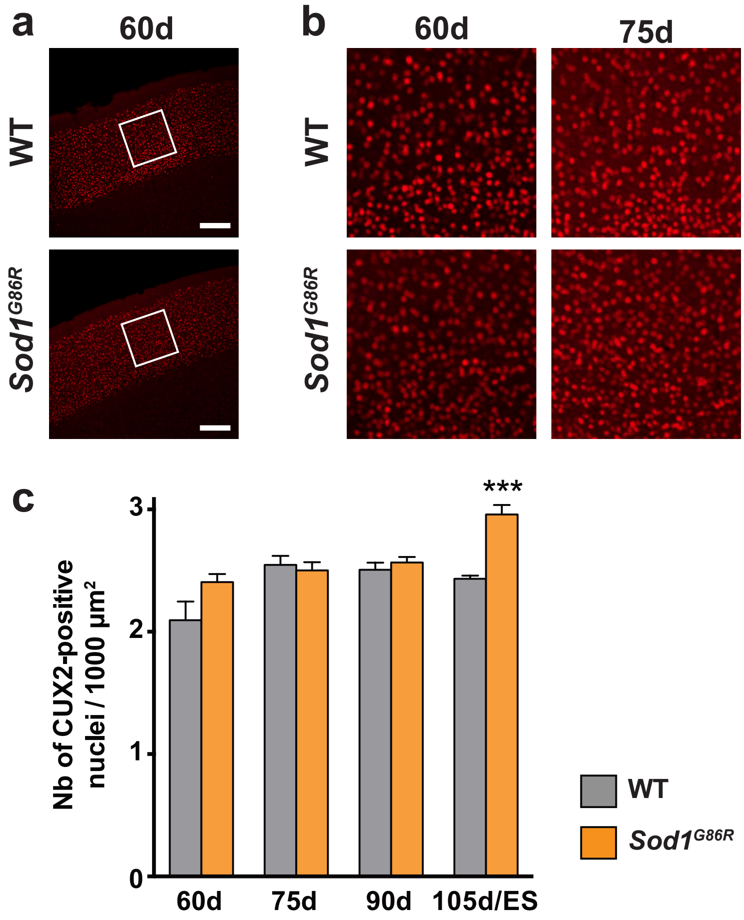

### Marques *et al.*, Supplementary Figure 6

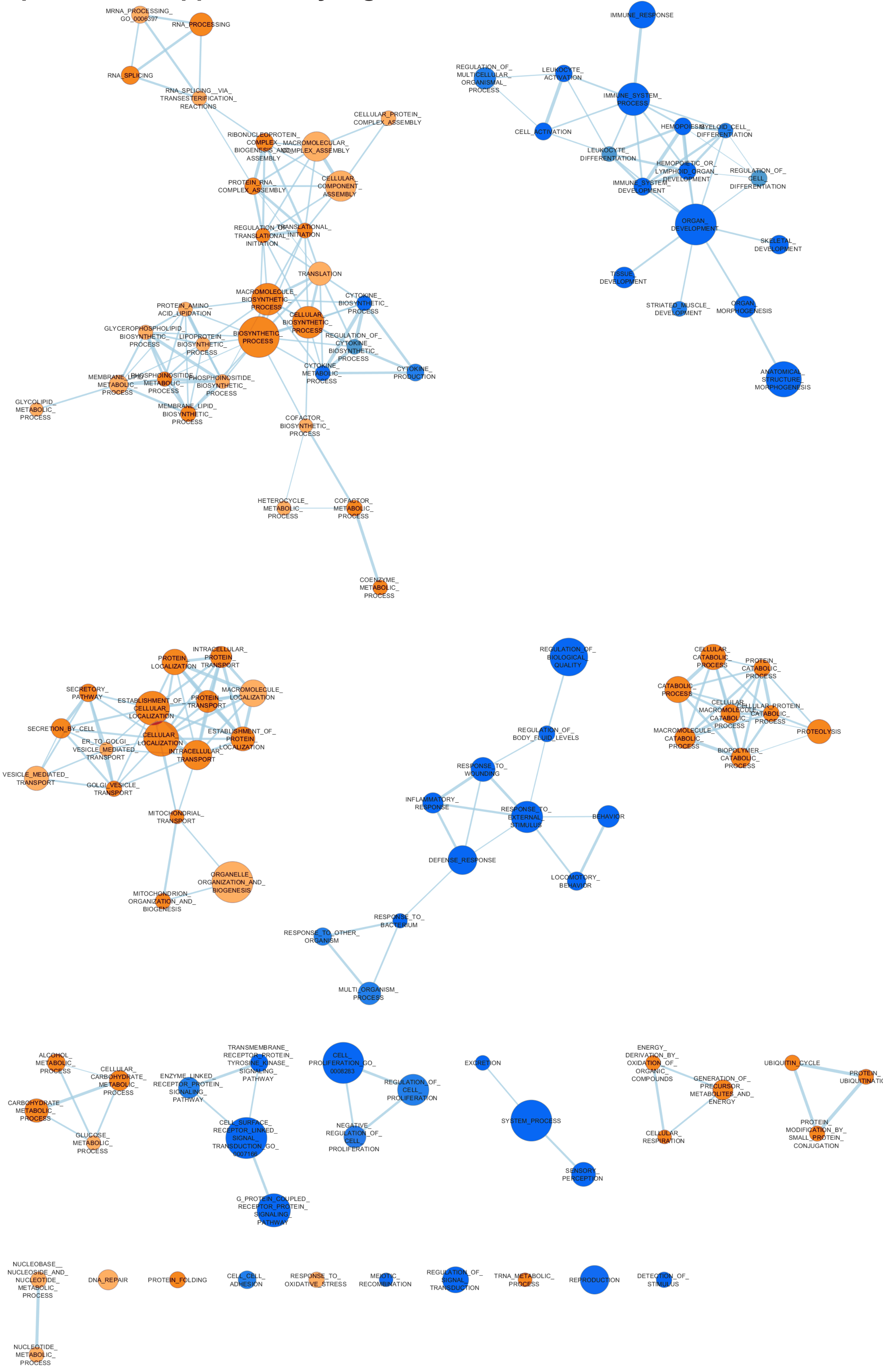

**Marques *et al.,* Supplementary Table 1: Gene set enrichment analysis (GSEA) of 29,017 ranked genes for GO terms of Cellular Component and Molecular Function.**

| GO Cellular Component | | | |
|---|---|---|---|
| NAME | NES | SIZE | FDR q-val |
| MITOCHONDRION | 6,93 | 304 | 0,000 |
| ORGANELLE_MEMBRANE | 5,72 | 263 | 0,000 |
| MITOCHONDRIAL_PART | 5,56 | 129 | 0,000 |
| ENVELOPE | 4,97 | 150 | 0,000 |
| ORGANELLE_ENVELOPE | 4,93 | 150 | 0,000 |
| MITOCHONDRIAL_MEMBRANE | 4,91 | 78 | 0,000 |
| MITOCHONDRIAL_ENVELOPE | 4,86 | 87 | 0,000 |
| MITOCHONDRIAL_INNER_MEMBRANE | 4,45 | 59 | 0,000 |
| MITOCHONDRIAL_MEMBRANE_PART | 4,43 | 47 | 0,000 |
| ORGANELLE_INNER_MEMBRANE | 4,34 | 67 | 0,000 |
| ENDOMEMBRANE_SYSTEM | 4,09 | 194 | 0,000 |
| RIBONUCLEOPROTEIN_COMPLEX | 3,60 | 117 | 0,000 |
| NUCLEAR_PART | 3,58 | 491 | 0,000 |
| PROTEASOME_COMPLEX | 3,41 | 22 | 0,000 |
| ENDOPLASMIC_RETICULUM | 3,21 | 244 | 0,000 |
| MEMBRANE_ENCLOSED_LUMEN | 3,20 | 395 | 0,000 |
| ORGANELLE_LUMEN | 3,20 | 395 | 0,000 |
| MITOCHONDRIAL_RESPIRATORY_CHAIN | 2,99 | 21 | 0,000 |
| MITOCHONDRIAL_MATRIX | 2,94 | 43 | 0,000 |
| MITOCHONDRIAL_LUMEN | 2,93 | 43 | 0,000 |
| ORGANELLAR_RIBOSOME | 2,82 | 21 | 0,000 |
| NUCLEAR_PORE | 2,80 | 26 | 0,000 |
| MITOCHONDRIAL_RIBOSOME | 2,79 | 21 | 0,000 |
| SPLICEOSOME | 2,66 | 40 | 0,000 |
| EXTRACELLULAR_REGION | -4,64 | 312 | 0,000 |
| EXTRACELLULAR_REGION_PART | -4,01 | 238 | 0,000 |
| EXTRACELLULAR_MATRIX | -3,58 | 83 | 0,000 |
| PROTEINACEOUS_EXTRACELLULAR_MATRIX | -3,45 | 82 | 0,000 |
| EXTRACELLULAR_SPACE | -3,18 | 160 | 0,000 |
| EXTRACELLULAR_MATRIX_PART | -2,86 | 53 | 0,000 |
| COLLAGEN | -2,71 | 21 | 0,000 |
| INTEGRIN_COMPLEX | -2,30 | 17 | 0,005 |
| BASEMENT_MEMBRANE | -2,17 | 36 | 0,010 |
| ACTIN_CYTOSKELETON | -2,08 | 117 | 0,015 |
| RECEPTOR_COMPLEX | -2,04 | 47 | 0,017 |

| GO Molecular Function | | | |
|---|---|---|---|
| NAME | NES | SIZE | FDR q-val |
| OXIDOREDUCTASE_ACTIVITY | 2,94 | 234 | 0,000 |
| ELECTRON_CARRIER_ACTIVITY | 2,66 | 69 | 0,003 |
| TRANSLATION_INITIATION_FACTOR_ACTIVITY | 2,64 | 20 | 0,003 |
| HYDROLASE_ACTIVITY__ACTING_ON_ACID_ANHYDRIDES | 2,60 | 204 | 0,003 |
| LIGASE_ACTIVITY | 2,57 | 92 | 0,003 |
| PYROPHOSPHATASE_ACTIVITY | 2,49 | 202 | 0,004 |
| GTPASE_ACTIVITY | 2,43 | 87 | 0,005 |
| UNFOLDED_PROTEIN_BINDING | 2,43 | 33 | 0,005 |
| TRANSLATION_REGULATOR_ACTIVITY | 2,38 | 33 | 0,006 |
| TRANSLATION_FACTOR_ACTIVITY__NUCLEIC_ACID_BINDING | 2,36 | 31 | 0,006 |
| OXIDOREDUCTASE_ACTIVITY__ACTING_ON_NADH_OR_NADPH | 2,35 | 22 | 0,006 |
| GENERAL_RNA_POLYMERASE_II_TRANSCRIPTION_FACTOR_ACTIVITY | 2,26 | 31 | 0,010 |
| NUCLEOSIDE_TRIPHOSPHATASE_ACTIVITY | 2,24 | 190 | 0,012 |
| SMALL_PROTEIN_CONJUGATING_ENZYME_ACTIVITY | 2,14 | 52 | 0,021 |
| SMALL_CONJUGATING_PROTEIN_LIGASE_ACTIVITY | 2,11 | 51 | 0,023 |
| PROTEIN_SERINE_THREONINE_PHOSPHATASE_ACTIVITY | 2,11 | 23 | 0,022 |
| INORGANIC_CATION_TRANSMEMBRANE_TRANSPORTER_ACTIVITY | 2,10 | 49 | 0,021 |
| SIGNAL_SEQUENCE_BINDING | 2,06 | 15 | 0,026 |
| GUANYL_NUCLEOTIDE_BINDING | 2,03 | 42 | 0,029 |
| UBIQUITIN_PROTEIN_LIGASE_ACTIVITY | 2,00 | 49 | 0,033 |
| HYDROGEN_ION_TRANSMEMBRANE_TRANSPORTER_ACTIVITY | 1,99 | 23 | 0,033 |
| ACID_AMINO_ACID_LIGASE_ACTIVITY | 1,94 | 56 | 0,042 |
| GTP_BINDING | 1,93 | 41 | 0,045 |
| RECEPTOR_ACTIVITY | -3,95 | 419 | 0,000 |
| TRANSMEMBRANE_RECEPTOR_ACTIVITY | -3,33 | 285 | 0,000 |
| TRANSMEMBRANE_RECEPTOR_PROTEIN_TYROSINE_KINASE_ACTIVITY | -3,02 | 38 | 0,000 |
| TRANSMEMBRANE_RECEPTOR_PROTEIN_KINASE_ACTIVITY | -2,91 | 46 | 0,000 |
| EXTRACELLULAR_MATRIX_STRUCTURAL_CONSTITUENT | -2,86 | 20 | 0,000 |
| PROTEIN_TYROSINE_KINASE_ACTIVITY | -2,68 | 55 | 0,001 |
| STRUCTURAL_MOLECULE_ACTIVITY | -2,68 | 185 | 0,001 |
| DNA_BINDING | -2,55 | 472 | 0,002 |
| SUBSTRATE_SPECIFIC_CHANNEL_ACTIVITY | -2,39 | 119 | 0,007 |
| ION_CHANNEL_ACTIVITY | -2,37 | 117 | 0,007 |
| PATTERN_BINDING | -2,33 | 38 | 0,009 |
| CATION_CHANNEL_ACTIVITY | -2,32 | 97 | 0,008 |
| TRANSCRIPTION_FACTOR_ACTIVITY | -2,25 | 264 | 0,011 |
| CARBOHYDRATE_BINDING | -2,23 | 55 | 0,012 |
| CYTOKINE_ACTIVITY | -2,15 | 69 | 0,016 |
| CHEMOKINE_RECEPTOR_BINDING | -2,15 | 27 | 0,016 |
| LIPASE_ACTIVITY | -2,15 | 39 | 0,015 |
| G_PROTEIN_COUPLED_RECEPTOR_ACTIVITY | -2,14 | 106 | 0,016 |
| GATED_CHANNEL_ACTIVITY | -2,08 | 99 | 0,021 |
| RECEPTOR_BINDING | -2,06 | 288 | 0,022 |
| CALCIUM_CHANNEL_ACTIVITY | -2,05 | 31 | 0,023 |
| G_PROTEIN_COUPLED_RECEPTOR_BINDING | -2,04 | 35 | 0,023 |
| GROWTH_FACTOR_BINDING | -2,03 | 26 | 0,023 |
| CHEMOKINE_ACTIVITY | -2,02 | 26 | 0,023 |

**Marques *et al*., Supplementary Table 2: Genes involved in RNA metabolism and previously linked to motoneuron and/or neurodevelopmental diseases.**

| Gene Symbol | Gene Name | Location | Related diseases *(Sources : Malacards.org: OMIN.org; PubMed.gouv)* |
|---|---|---|---|
| ***Aars*** | alanyl-tRNA synthetase | 16q22.1 | Charcot-Marie-Tooth disease, axonal, types 2e and 2n; Epileptic encephalopathy, early infantile, 29 |
| ***Dars*** | Aspartyl-tRNA synthetase | 2q21.3 | Hypomyelination with brainstem and spinal cord involvement and leg spasticity |
| ***Ddx20*** | DEAD (Asp-Glu-Ala-Asp) box polypeptide 20 | 1q13.2 | Spinal Muscular Atrophy |
| ***Eif2b2*** | Eukaryotic Translation Initiation Factor 2B Subunit Beta | 14q24.3 | Leukoencephalopathy with vanishing white matter; Ovarioleukodystrophy |
| ***Eif2b3*** | Eukaryotic Translation Initiation Factor 2B Subunit Gamma | 1p34.1 | Leukoencephalopathy with vanishing white matter; Leukodystrophy |
| ***Eif2b4*** | Eukaryotic Translation Initiation Factor 2B Subunit Delta | 2q23.3 | Leukoencephalopathy with vanishing white matter; Ovarioleukodystrophy |
| ***Eif2b5*** | Eukaryotic Translation Initiation Factor 2B Subunit Epsilon | 3q27.1 | Leukoencephalopathy with vanishing white matter; Ovarioleukodystrophy |
| ***Fars2*** | Phenylalanyl-TRNA Synthetase 2, Mitochondrial | 6p25.1 | Infantile-onset *FARS2* definicency with epileptic encephalopathy and lactic acidosis; Spastic paraplegia, autosomal recessive |
| ***Gemin8*** | Gem Nuclear Organelle Associated Protein 8 | Xp22.2 | Spinal Muscular Atrophy |
| ***Kars*** | Lysyl-tRNA synthetase | 16q23.1 | Charcot-Marie-Tooth disease; Deafness |
| ***Nono*** | Non-POU d-Containing Octamer Binding Prot. Phos-ase 1, Reg. SU 114 | Xq13.1 | X-linked syndromic intellectual disability |
| ***Nsun2*** | NOP2/Sun RNA Methyltransferase Family Member 2 | 5p15.31 | Dubowitz syndrome; Autosomal recessive non-syndromic intellectual disability |
| ***Prpf31*** | Pre-MRNA Processing Factor 31 | 19q13.42 | Retinis Pigmentosa 11 |
| ***Rars*** | Arginyl-tRNA Synthetase | 5q34 | Leukodystrophy hypomyelinating |
| ***Sars*** | Seryl-tRNA Synthetase | 1q13.3 | Autosomal recessive intellectual disability |
| ***Snrpb*** | Small Nuclear Ribonucleoprotein Polypeptides B And B1 | 20q13 | Cerebrocostomandibular syndrome |
| ***Snrpn*** | Small Nuclear Ribonucleoprotein Polypeptide N | 15q11.2 | Prader-Willi Syndrom; Angelman Syndrome; Autism |
| ***Trnt1*** | tRNA Nucleotidyl Transferase 1 | 3p26.2 | Retinitis pigmentosa with erythrocytic microcytosis; Sideroblastic anemia with B-cell immunodeficiency, periodic fevers, and developmental delay; Mental retardation |
| ***Wars*** | Tryptophanyl-tRNA Synthetase | 14q32.2 | Growth retardation and progressive leukoencephalopathy |
| ***Yars*** | Tyrosyl-tRNA Synthetase | 1p35.1 | Charcot-Marie-Tooth disease |
| ***Eftud2*** | Elongation Factor Tu GTP Binding Domain Containing 2 | 17q21.31 | Mandibulofacial dysostosis with microcephaly |
| ***Sf3b4*** | Splicing Factor 3b Subunit 4 | 1q21.2 | Nager syndrome |
| ***Slbp*** | Stem-Loop Binding Protein | 4q16.3 | Wolf-Hirschhorn Syndrome |

Marques *et al.*, Supplementary Table 3: Gene set enrichment analysis (GSEA) of 1,163 ranked mis-spliced genes, for the GO terms Biological Process, Cellular Component and Molecular Function.

| GO Biological Process | | | |
|---|---|---|---|
| NAME | NES | SIZE | p value |
| POSITIVE_REGULATION_OF_INTRACELLULAR_TRANSPORT | 2,0 | 16 | 0,001 |
| SYNAPTIC_SIGNALING | 1,8 | 28 | 0,005 |
| REGULATION_OF_NEURON_DIFFERENTIATION | 1,8 | 33 | 0,002 |
| REGULATION_OF_CYTOPLASMIC_TRANSPORT | 1,8 | 23 | 0,002 |
| SMALL_MOLECULE_CATABOLIC_PROCESS | 1,8 | 18 | 0,004 |
| DEPHOSPHORYLATION | 1,8 | 16 | 0,010 |
| POSITIVE_REGULATION_OF_PHOSPHORUS_METABOLIC_PROCESS | 1,8 | 53 | 0,001 |
| POSITIVE_REGULATION_OF_MAPK_CASCADE | 1,8 | 22 | 0,010 |
| REGULATION_OF_NEURON_PROJECTION_DEVELOPMENT | 1,8 | 32 | 0,005 |
| REGULATION_OF_NEURON_DEATH | 1,8 | 19 | 0,012 |
| HEART_DEVELOPMENT | 1,8 | 29 | 0,008 |
| RNA_CATABOLIC_PROCESS | 1,7 | 20 | 0,018 |
| RESPONSE_TO_CYTOKINE | 1,7 | 30 | 0,014 |
| REGULATION_OF_KINASE_ACTIVITY | 1,7 | 47 | 0,006 |
| NEURON_DEVELOPMENT | 1,7 | 45 | 0,003 |
| LOCOMOTORY_BEHAVIOR | 1,7 | 18 | 0,020 |
| REGULATION_OF_INTRACELLULAR_PROTEIN_TRANSPORT | 1,7 | 20 | 0,013 |
| SINGLE_ORGANISM_CATABOLIC_PROCESS | 1,7 | 58 | 0,005 |
| NEURON_DIFFERENTIATION | 1,7 | 50 | 0,014 |
| REGULATION_OF_NERVOUS_SYSTEM_DEVELOPMENT | 1,7 | 39 | 0,012 |

| GO Cellular Component | | | |
|---|---|---|---|
| NAME | NES | SIZE | p value |
| CYTOSOLIC_PART | 2,1 | 15 | 0,000 |
| AXON_PART | 1,7 | 20 | 0,014 |
| AXON | 1,6 | 34 | 0,027 |
| SECRETORY_VESICLE | 1,5 | 27 | 0,050 |
| CATALYTIC_COMPLEX | 1,5 | 96 | 0,019 |
| INTRINSIC_COMPONENT_OF_PLASMA_MEMBRANE | 1,5 | 53 | 0,050 |

| GO Molecular Function | | | |
|---|---|---|---|
| NAME | NES | SIZE | p value |
| RECEPTOR_BINDING | 1,8 | 64 | 0,003 |
| PROTEIN_KINASE_ACTIVITY | 1,7 | 41 | 0,011 |
| RECEPTOR_ACTIVITY | 1,7 | 30 | 0,009 |
| SIGNALING_RECEPTOR_ACTIVITY | 1,7 | 24 | 0,017 |
| GATED_CHANNEL_ACTIVITY | 1,6 | 15 | 0,025 |
| KINASE_BINDING | 1,6 | 32 | 0,016 |
| KINASE_ACTIVITY | 1,6 | 55 | 0,011 |
| TRANSFERASE_ACTIVITY_TRANSFERRING_PHOSPHORUS_CONTAINING_GROUPS | 1,6 | 68 | 0,016 |
| PASSIVE_TRANSMEMBRANE_TRANSPORTER_ACTIVITY | 1,6 | 19 | 0,041 |
| PHOSPHORIC_ESTER_HYDROLASE_ACTIVITY | 1,6 | 17 | 0,034 |
| UBIQUITIN_LIKE_PROTEIN_LIGASE_BINDING | 1,5 | 24 | 0,045 |
| PROTEIN_DIMERIZATION_ACTIVITY | 1,5 | 64 | 0,031 |
| TRANSFERASE_ACTIVITY_TRANSFERRING_ACYL_GROUPS | -1,7 | 20 | 0,000 |
| TRANSFERASE_ACTIVITY_TRANSFERRING_ACYL_GROUPS_OTHER_THAN_AMINO_ | -1,7 | 19 | 0,013 |
